# Lateral hypothalamus directs stress-induced modulation of acute and psoriatic itch

**DOI:** 10.1101/2025.06.09.658592

**Authors:** Jagat Narayan Prajapati, Aynal Hoque, Manojeet Pattanayak, Giriraj Sahu, Arnab Barik

**Affiliations:** Centre for Neuroscience, Indian Institute of Science, Bengaluru, Karnataka, India- 560012; Molecular Biophysics Unit, Indian Institute of Science, Bengaluru, Karnataka, India- 560012; Centre for High Impact Neuroscience and Translational Applications (CHINTA), TCG CREST, Kolkata, India

## Abstract

Stress and anxiety are well-known modulators of both physiological and pathological itch. Acute stress suppresses itch, while chronic stress exacerbates it. These effects are mediated by neural circuits within the brain, though the precise mechanisms remain poorly understood. In this study, we investigate the role of neurons in the stress-sensitive lateral hypothalamic area (LHA) in modulating itch. Using neural activity-dependent genetic labeling and chemogenetic tools, we selectively engaged a population of LHA neurons (LHA^stress-TRAP^ neurons) responsive to stress. Transient stimulation of these neurons induced anxiety-like behaviors, conditioned place aversion, and suppressed acute (chloroquine-induced) and chronic (psoriatic) itch. Conversely, the inhibition of the LHA^stress-TRAP^ neurons enhanced acute and chronic itch. Interestingly, LHA^stress-TRAP^ neurons did not respond to acute itch stimuli, but their activity was temporally correlated with scratching episodes in mice with psoriasis. Ex vivo whole-cell patch-clamp recordings revealed that these neurons exhibit heightened excitability in psoriatic animals. Anterograde viral tracing demonstrated that LHA^stress-TRAP^ neurons project to brainstem regions implicated in itch modulation, including the periaqueductal gray (PAG), rostral ventromedial medulla (RVM), and lateral parabrachial nucleus (LPBN). Furthermore, chemogenetic activation and optogenetic silencing of LHA^stress-TRAP^ axon terminals revealed that bidirectional modulation of itch is primarily mediated through projections to the PAG. Together, these findings identify a previously unrecognized central mechanism by which stress modulates itch, centered on a specific population of LHA neurons and their downstream brainstem targets.

## Introduction

Itch and pain are distinct behavioral responses to noxious somatosensory stimuli: pruritic stimuli, including allergens, insect bites, and woolen textiles, elicit itch, while algesic stimuli such as noxious heat, cold, or mechanical insult evoke pain. Despite their sensory specificity, both modalities are modulated by internal brain states, including chronic stress and anxiety ^1–3^. While recent studies have elucidated key neural circuits through which stress interacts with pain perception and processing^4–6^, the mechanisms by which stress modulates itch and how these systems may intersect remain poorly understood.

In animal models of acute stress caused by physical restraint or forced swimming, behavioral responses to algesic and pruritic chemical stimuli are suppressed ^7^. Interestingly, the extent of itch modulation was proportional to the severity of stress. Similarly, acute stress attenuated mechanically (cowhage) evoked itch in human subjects ^8^. Despite these observations, the neural circuit mechanisms underlying stress-induced modulation of itch remain poorly understood. Traditionally, it was thought that the central and basolateral amygdala are the primary players in stress modulation of itch ^9,10^. However, it is not clear if other brain areas play a role in the interactions between stress and itch. The lateral hypothalamic area (LHA), usually associated with energy homeostasis, motivated behaviors, and arousal^11–13^, has been shown to drive stress and anxiety ^14–18^. Early lesion studies demonstrated that damage to the LHA results in diminished responsiveness to somatosensory stimuli^19^. Recently, it was shown that the excitatory LHA neurons receiving input from the anxiogenic lateral septum (LS) mediated stress-induced analgesia by suppressing pro-nociceptive neurons in the rostral ventromedial medulla (RVM)^4^. Conversely, a distinct population of inhibitory LHA neurons downstream of the LS mediated chronic itch-induced anxiety^20^. The thalamic reuniens nucleus (Re), which lies upstream of the LS-LHA circuitry, was shown to transform pruritic sensory input into stress and anxiety^20^.

However, whether and how LHA neurons contribute specifically to the modulation of itch by acute stress remains unclear. Several brain regions that receive noxious somatosensory input and are downstream of the LHA, including the lateral parabrachial nucleus (LPBN), periaqueductal gray (PAG), and RVM, are known to regulate pruritogen-induced scratching and itch-related aversive behaviors. For instance, silencing LPBN neurons eliminates chloroquine-induced scratching^21–23^, while excitatory and inhibitory neurons in the ventrolateral PAG facilitate and inhibit itch^24–26^, respectively. RVM neurons modulate pruriceptive responses through their synaptic inputs to the spinal cord^27,28^.

Here, we investigated whether neurons in the lateral hypothalamic area (LHA) mediate stress-dependent modulation of itch. To this end, we utilized the TRAP2 transgenic mouse line ^29,30^, in which tamoxifen-inducible CreERT2 is expressed under the control of the immediate early gene *cFos* promoter, enabling activity-dependent and temporally precise genetic access to neurons via 4-hydroxytamoxifen (4-OHT) administration. By combining TRAP2 mice with locally delivered Cre-dependent AAV vectors, we selectively targeted the stress-activated LHA neurons (LH^Stress-TRAP^) to assess their role in chloroquine-induced acute itch and imiquimod-induced chronic psoriatic itch. Using temporally restricted chemogenetic activation and inactivation strategies, we tested whether LH^Stress-TRAP^ neurons are sufficient and necessary for itch modulation. In vivo, calcium imaging and ex vivo electrophysiological recordings were employed to characterize the sensory and stress-related stimuli that recruit and alter the activity of these neurons. Additionally, anterograde and retrograde tracing approaches defined the connectivity and projection targets of LH^Stress-TRAP^ neurons. Together, these circuit-level analyses demonstrate that stress-responsive LHA neurons play a key role in regulating both physiological and pathological itch.

## Methods

### Mouse line

Animal care and experimental procedures were performed following protocols approved by the CPSCEA at the Indian Institute of Science. TRAP2 (Fos^2A-iCreERT2^)^29^ mice, stock number 030323, were purchased from Jackson Laboratory. The animals were housed at the Central Animal Facility (CAF) under standard transgenic animal housing conditions in a 12-hour light-dark cycle with ad libitum access to food and water. Genotyping was performed according to the protocols of Jackson Laboratories. CD-1 mice (7-12 weeks old) were bought from CAF for the behavioural experiments. An equal number of males and females were used in all the behavioural experiments. All mice used in the behavioral assays were between 7 and 12 weeks old. All the behaviors were done during the light cycle.

### Viral Vectors

Vector used and sources: pAAV5-hsyn-DIO-EGFP (Addgene, Catalog# v78581-AAV9, titer-2.5 x 10^13^ GC/ml), pAAV5-FLEX-tdTomato (Addgene, Catalog# 28306-AAV1, titer-1.6 x 10^13^ GC/ml), pAAV5-hsyn-DIO-hM3D(Gq)-mCherry (Addgene, Catalog# 44361, titer-1.8 x 10^13^ GC/ml), rAAV2/9-EF1α-DIO-Kir2.1-P2A-EGFP (BrainVTA, Catalog# PT-1401, titer-2 x 10^12^ vg/ml), AAV9.syn.flex.GcaMP8s (Addgene, Catalog# 162377, titer-2.7 x 10^13^ GC/ml), rAAV5-EF1α-DIO-oRVG (BrainVTA, Catalog# PT-0023, titer-2 x 10^12^ vg/ml), rAAV5-EF1a-DIO-H2B-eGFP-T2A-TVA (BrainVTA, Catalog# PT-0021, titer-2 x 10^12^ vg/ml), and RV-CS-N2C-deltaG-tdTomato (BrainVTA, Catalog# R05002, titer-2 x 10^8^ vg/ml), scAAV-1/2-hSyn1-FLPO-SV40p(A) (University of Zurich, Catalog# v59-1, titer-6.7 x 10^12^ vg/ml), pAAV-Ef1a-fDIO-tdTomato (Addgene, Catalog# 128434-AAV1, titer-1.8 x 10^13^ GC/ml).

### Antibodies

Chicken anti-GFP antibody (aveslabs catalog# 1010), Phospho-c-Fos (Ser32) Rabbit monoclonal antibody (Cell Signaling Technology Catalog# 5348), Goat anti-tdTomato (SICGEN catalog# AB8181), Goat anti-Chicken IgY (H+L) secondary Antibody, Alexa Fluor™ 488 (Invitrogen catalog# A11039), Donkey anti-Rabbit IgG (H+L) secondary Antibody, Alexa Fluor™ 488 (Invitrogen catalog# A21206), Donkey anti-Goat IgG (H+L) secondary Antibody, Alexa Fluor™ 594 (Invitrogen catalog# A11058),

### Stereotaxic injections

Mice were anesthetized with 2 % isoflurane/oxygen before and during the surgery and mounted on the stereotaxic frame (RWD 69100 Rotational Digital Stereotaxic Frame). An incision was made to expose the skull, and subsequently, the skull was aligned to the horizontal plane. Craniotomy was performed at the marked point using a hand-held micro-drill (RWD). A Hamilton syringe (10 ul) with a glass pulled needle was used to infuse 300 nL of viral particles (1:1 in saline) at a 100 nL/min rate. The following coordinates introduced the virus: LH-AP: −1.70, ML: ±1.00, DV: −5.15. For rabies tracing experiments, rAAV5-EF1α-DIO-oRVG and rAAV5-EF1α-DIO-EGFP-T2A-TVA were injected first, followed by RV-CS-N2C-deltaG-tdTomato one week after stressTRAPing. Tissue was harvested after 1 week of rabies injection for histochemical analysis. The study did not include the brain tissues where apparent cell death was observed through morphological examination. Post-hoc histological examination of each injected mouse was used to confirm that viral-mediated expression was restricted to the target nuclei.

### Fiber optic cannula and stainless steel guide cannula implantation

Fiber optic cannula from RWD (Ø1.25 mm Ceramic Ferrule, 300 μm Core, 0.39NA, L = 7 mm, catalog# R-FOC-BL300C-39NA) was implanted at AP: −1.70, ML: +1.00, DV: −5.15 in the LH of the AAV-DIO-GCaMP8s infused mice. The cannula was fixed to the skull using light-cured dental cement (GC corporation powder-catalog# 002505, liquid-catalog# 002524). Animals were allowed to recover for at least 1 week before performing behavioral tests. Successful labeling and fiber implantation were confirmed post hoc by staining for GFP for viral expression and injury caused by the fiber, respectively. Only animals with viral-mediated gene expression and fiber implantations at the intended locations, as observed in post hoc tests, were included in the analysis.

For chemogenetic activation of LHA^stress-TRAP^ neuron terminal a stainless steel guide cannula (O.D. 0.48mm, 26G, 5mm, RWD catalog# 62003) were implanted bilaterally at the PAG (AP: −4.4, ML: ±1.23, DV: - 2.86, α=15°) and PBN (AP: −5.34, ML: 1.00, DV: 3.15) in the stress TRAPed mice. For RVM terminals, an activation 8mm cannula (O.D. 0.48mm, 26G) was implanted at AP: −5.8, ML: 0.10, DV: −5.50 into the RVM of the stress TRAPed mice. The cannulas were fixed to the skull using dental cement, and the animals were allowed to recover for a week before performing behavioral tests. Mice were lightly anesthetized using isoflurane, and the injection tube was inserted into the guide cannula. The infusion cannula was connected to the microinfusion pump (KD Scientific, catalog# 78-8130). Saline/DCZ was infused at the rate of 100 nl/min. The infusion cannula was removed 10 minutes after the infusion, and 15 minutes later, behavioural assays were done. Successful implantations were confirmed post hoc by staining for Fos at the intended location, along with viral expression and injury caused by the cannula. Only animals with viral-mediated gene expression and cannula implantations at the intended locations, as observed in post hoc tests, were included in the analysis.

### Stress TRAPing

4-hydroxytamoxifen (4-OHT; Hello Bio, UK, Cat No.. H6040) was prepared by dissolving it in ethanol at a 20 mg/ml concentration^29^. The solution was aliquoted and stored at −40°C for several days. 4-OHT was redissolved just before use and mixed with corn oil in a 1:1 ratio. 4-OHT (50 mg/kg body weight) was intraperitoneally administered to the mice, and 15 minutes later, the mice were subjected to 1-hour restraint stress by placing the mouse in a 50 ml Falcon tube, which has holes for proper air ventilation. All the behavioral and anatomical studies were done one week after the stress TRAPing.

### Fiber photometry

A dual-channel fiber photometry system from RWD (R810) was used to collect the data^31,32^. The light from two light LEDs (410 and 470 nm) was passed through a fiber optic cable (RWD-Ø1.25 mm Ceramic Ferrule, 200 μm Core, 0.39NA, L = 2 mm, catalog# R-FC-L-N3-200-L1) coupled to the cannula implanted in the mouse. Fluorescence emission was acquired through the same fiber optic cable onto a CMOS camera through a dichroic filter. Mice were lightly anesthetized, and the fiber-optic cable was connected to the optical cannula attached to the mouse skull. Mice were habituated to the fibers for 2 days before performing any behavioural assays. The output power was adjusted to 30%, which gives 20-50 µW power at the fiber tip. The signals were acquired at a 30 fps frame rate. The data was analyzed using the RWD photometry software, and .csv files were generated. The start and end of stimuli were timestamped. All trace graphs were plotted from .csv files using GraphPad Prism software version 8.

### Chemogenetic activation

For chemogenetic activation of LHA^stress-TRAP^ neurons, deschloroclozapine (DCZ) (Hello Bio, catalog# HB9126), 2 µg/kg body weight, was administered intraperitoneally (i.p.) into the stress-TRAPed mice ^33^. All the behavioral assays were done 15 min after the DCZ administration.

### Optogenetic silencing

For optogenetic inhibition of PAG terminals of the LHA^stress-TRAP^ neurons using halorhodopsin eNPHR3.0^34^, a bilateral fiber optic cannula (Ø1.25 mm Ceramic Ferrule, 200 μm Core, 0.22NA, L = 5 mm, catalog# R-FOC-L200C-22NA) was implanted at PAG (AP: −4.4, ML: ±1.23, DV: −2.86, α=15°). One week after the implantation, mice were habituated for 2 days and then used for behavioural experiments. Prizmatix Optogenetics-LED-Yellow was used to deliver a 595 nm constant light to inhibit the PAG terminals of the LHA^stress-TRAP^ neurons. Mice were briefly anesthetized using isoflurane, and an optical fiber (Prizmatix, L = 2m, core diameter 500 µm, and NA 0.63) was connected to the fiber optic cannula to deliver light. The scratching behaviour of mice is recorded with a 5-minute light ON and a 5-minute light OFF cycle for 1 hour.

### Brain slice preparation and electrophysiology

TRAP2 mice were injected with AAV encoding Cre-dependent eGFP and, 3 weeks later, stress-TRAPed. One week later, control mice were treated with daily topical application of moisturizer cream (PONDS), while the chronic mice received daily application of imiquimod for 6 days. Control and psoriatic mice were anesthetized with 4% isoflurane, followed by decapitation and surgical dissection of the brain. Coronal brain slices of 300 μm were prepared using semi-automated vibratome (VT1200S; Leica Microsystems, Germany) in ice-cold cutting solutions composed of in mM: sucrose (75), NaCl (87), NaHCO₃ (25), Na₂HPO₄ (1.25), CaCl₂ (0.5), and MgCl₂ (7) continuously perfused with carbogen (5% CO2 + 95% O2) gas. The brain slices were placed in a 32 °C water bath for 15 minutes followed by incubation at room temperature (∼25 °C) for at least an hour in artificial cerebrospinal fluid (ACSF) containing in mM: NaCl (126), KCl (2.5), NaHCO₃ (25), Na₂HPO₄ (1.25), CaCl₂ (1.5), MgCl₂ (1.5), CaCl₂ (1.5), and glucose (25) under constant perfusion with carbogen gas, before being considered for patch-clamp experiments. During recording, the brain slices were shifted to a recording chamber bathed with ACSF under carbogen perfusion and maintained at 32 °C by using a digitized temperature controller (Warner Instruments, USA).

For whole-cell recordings, the LH neurons were identified under 40X magnification, displaying eGFP fluorescence emission by using a dot-contrast enabled IR-DIC compatible upright patch clamp microscope (Axio Examiner D1, Carl Zeiss, Germany). Thick-walled borosilicate glass capillaries (OD: 1.5 mm, ID: 0.86 mm) were used for preparing the patch pipettes having 3-5 mΩ resistances by using a horizontal micropipette puller (Sutter Instruments, USA). The patch pipettes were filled with an internal solution composed of in mM: K-gluconate (135), KCl (4), Na2ATP (5), NaGTP (0.5), HEPES (10), adjusted pH to 7.3 with KOH. Whole-cell current clamp experiments were performed using a computer-controlled Multiclamp 700B amplifier operated through pClamp 11.3 software (Molecular Devices, USA). The current and voltage traces were low-pass filtered with 2 kHz and digitized at 10 kHz using a hum-silencer-enabled Digidata 1550B (Molecular Devices, USA). The resting membrane potential was noted immediately after attaining whole-cell mode without any external current injection. Postsynaptic step current injections of 3 seconds, ranging from 0 to +200 pA with 20 pA current increments, were used to assess the gain of firing rate (F/A plot) of LHA neurons. Similarly, for determining the input resistance, −100 to +100 pA postsynaptic current was injected through a patch pipette with a 10-pA increment. All the electrophysiological data were analyzed using the Clampfit module of pClamp software, and Adobe Illustrator was used to plot the graphs.

### Behavioural assays

#### Chloroquine-induced itch assay

The nape of the neck of mice was shaved with a hand-held Philips shaver 2–3 days before behavioral experimentation, and the mice were habituated in the behavior room. Unless otherwise stated, the mice used for behavioral studies were blinded prior to initiation of the studies by an individual not involved in the experimentations described here. All behavioral experiments were quantified by one experimenter and randomly cross-verified by another. All itch experiments were videotaped with a Logitech camera, and videos were acquired through vendor-supplied software. Mice were individually placed in four-part plexiglass chambers with chamber dimensions of 6 cm X 6 cm X 14 cm. The roof of the chamber had holes for air ventilation. Animals were habituated in the chamber for 15 min before chloroquine injections. DCZ 2 µg/kg body weight was administered intraperitoneally (i.p.) 15 min before chloroquine injection. Chloroquine (375 µg/75 µl) (Sigma Catalog# C6628) was administered intradermally into the nape of the neck of the mice, and the subsequent scratching behavior was recorded for 30 min^35^. Hind leg-directed scratching of the nape was characterized as a scratch, and the videos were quantified, blinded to the experimental conditions.

#### Imiquimod induced psoriatic itch

Imiquimod (5% w/w from Glenmark,) was used to induce psoriasis in mice ^36,37^. To induce psoriasis-like chronic itch, the nape of each mouse was shaved using Veet hair removal cream, and imiquimod was topically applied once daily to the shaved area for six consecutive days. This treatment reliably induced inflamed, scaly skin lesions characterized by thickened and dry epidermis. Mice were manually inspected, and only those exhibiting pronounced psoriatic features—namely, inflamed, thickened, and scaly skin—were selected for behavioral analysis. Following psoriasis induction, mice were individually placed in four-compartment plexiglass chambers for habituation, after which their behavior was recorded for 30 minutes. Spontaneous scratching was defined as hind limb-directed contact to the nape region. Scratching bouts were quantified by observers blinded to the experimental conditions.

#### Hotplate test

The thermal hotplate experiments were performed using the Hot and Cold Plate analgesiometer (HC-01, Orchid Scientific)^38,39^. The specifications of the instrument used are-enclosure size: 205 × 205 × 250 mm; plate size: 190 × 190 × 06 mm; temperature range: −5°C to 60 °C. A single experimenter introduced the mice into the enclosure on the thermal plate across all the experiments and performed analysis in a blinded manner. The mice were habituated in the experimental room for 30 minutes and in the enclosure for 5 minutes at 32 °C before the experimentation for three consecutive days. On the experimental day, mice were placed on the hotplate at 52 °C, and the behavior was recorded for 45 seconds using three Logitech web cameras placed at the left, right, and front angles around the hotplate^38^. Later, videos were quantified individually for any nocifensive behaviors (licks, shakes, and jumps) exhibited by the mice, blinded to the experimental conditions.

#### Open field test

The open field test was used to evaluate anxiety-like behaviour^40^. The open field arena was made up of acrylic white opaque walls with dimensions 50 cm (length) x 50 cm (width) x 38 cm (height). The central field has a dimension of 30 cm x 30 cm. The mouse was placed in the middle of the arena and allowed to move freely for 10 min. The movement was recorded using an overhead-mounted Logitech camera. The open field arena was cleaned with 70% alcohol between every trial. The total distance moved and the time spent in the central field was tracked using DeepLabCut.

#### Light-Dark Box test

The light-dark box with dimensions 40 cm (length) x 20 cm (width) x 36 cm (height) of each chamber (light and dark chamber) was used to measure the anxiety-like behavior^41,42^. Mice could freely access both the light and dark chambers via a small opening which connects both chambers. The mice were introduced into the light side of the apparatus and allowed to explore freely for 15 min, and the movement was recorded using a Logitech camera (C930e) from the top. The time spent in the light box is taken as the degree of anxiety in the mouse. The apparatus was cleaned with 70% alcohol between every trial. The movement of the mouse was tracked and plotted using DeepLabCut.

#### Conditioned Place Aversion (CPA) test

A three-compartment custom-built CPA apparatus was used to test the conditioned place aversion in mice. Both outer chambers have a dimension of 32 cm (length) x 32 cm (width) x 28 cm (height), while the middle chamber has a dimension of 11 cm x 8.5 cm. One of the outer chambers has white stripe walls and a steel mesh floor; the other has black walls and a steel rod floor; the middle has gray walls and a smooth PVC floor as a neutral zone. Two manual doors between these three chambers can be closed to block entry into any of the chambers. The light intensity was constant to prevent any innate preference. Each chamber was cleaned thoroughly with 70% ethanol between every trial. The mouse movement was videotaped using a Logitech camera (C930e).

The unbiased CPA experiment ends in 5 days^43^. On Day 1 (pre-conditioning phase), mice were placed in the central compartment of a three-chamber apparatus and allowed to explore all chambers for 15 minutes. The time spent in each outer chamber (T1 and T2) was recorded. Mice exhibiting a preference ratio (T1/T2) between 2:3 and 3:2 were selected for further experiments; animals outside this range were excluded to ensure unbiased baseline preferences. From Days 2 to 4 (conditioning phase), animals underwent two daily conditioning sessions (morning and evening), separated by a minimum interval of four hours. On Day 2, the chamber with striped walls was designated as the drug-paired environment (DCZ), while the chamber with black walls was designated as the saline-paired environment. In the morning session, mice received an intraperitoneal (i.p.) injection of DCZ and were confined to the drug-paired chamber for 30 minutes. In the evening session, they received an i.p. injection of saline and were confined to the saline-paired chamber for 30 minutes. On Day 3, the conditioning sequence was reversed: mice received saline in the morning and DCZ in the evening, with corresponding chamber confinement. On Day 4, the original sequence was reinstated, with DCZ administered in the morning and saline in the evening. On Day 5 (post-conditioning phase), both chamber doors were opened, and mice were allowed to explore the entire apparatus freely for 15 minutes. Time spent in the DCZ-paired chamber was compared between Day 1 and Day 5 to evaluate the development of conditioned place aversion (CPA).

#### DeepLabCut for tracking mice

The tracking of mice in the open field test, light-dark box test, and CPA test were done using DeepLabCut (DLC) (version 2.3.9)^38,44^. Data were processed and analyzed on a custom-built workstation equipped with an AMD Ryzen 9 5900X 12-core processor and an NVIDIA GPU. For training the DeepLabCut (DLC) model, 20 frames were manually labeled from each of the five videos. Following training, the model was used to analyze the videos, generating position plots and corresponding output in .csv format. For visualization, representative plots of tracked spine positions were produced.

#### Immunostaining, multiplex in situ hybridization, and confocal microscopy

Mice were anesthetized with isoflurane and perfused transcardially with 1X Phosphate Buffered Saline (PBS) (Takara catalog# T9181) and 4 % Paraformaldehyde (PFA) (Ted Pella, Inc. catalog# 18505), harvested brains and spinal cords were further fixed in 4 % PFA, overnight, and subsequently transferred to 15 % and 30 % sucrose for serial dehydration. Brain tissues were placed in the Cryo-Embedding Compound (Ted Pella, Inc.) and frozen at −40°C. Subsequently, 50 µm-thick coronal brain sections were cut using a cryostat (RWD Minux FS800). For immunostaining experiments, tissue sections were rinsed in 1X PBS (3 times) and incubated in the blocking buffer (5 % Bovine Serum Albumin (BSA) + 0.5 % Triton X-100 + 1X PBS) (BSA-HIMEDIA catalog# TC194, Triton X-100 SRL catalog# 64518) for one hr at room temperature. Sections were then incubated in the primary antibody (dilution 1:1000 X in blocking buffer) at room temperature overnight. Sections were rinsed 3 times with 1X PBS + 0.5 % Triton X-100 solution and incubated for two hr in Alexa Fluor conjugated goat anti-rabbit/ chicken or donkey anti-goat/rabbit secondary antibodies (dilution 1:1000 X in blocking buffer) along with DAPI (SRL catalog# 18668) at room temperature. Then sections were washed with 1X PBS + 0.5 % Triton X-100, and mounted onto charged glass slides (Ted Pella, Inc. catalog# 260382-3). Citifluor AF-1 mounting media (Ted Pella, Inc. catalog# 19470-1) was used to cover-slip (Blue star microscopic cover glass 24 x 60 mm 10 Gms) the slides. Subsequently, sections were imaged on the upright fluorescence microscope (Khush Enterprises, Bengaluru) (2X, 4X, and 10X lenses) and a Confocal Microscope (Leica SP8 Falcon, Germany). ImageJ/FIJI processing software was used to process the images. Confocal images were processed using the Leica image analysis suite software.

Fresh brains were rapidly harvested and flash-frozen at –80 °C for subsequent in situ hybridization (ISH). Coronal sections (20 µm) were prepared using a cryostat. Multiplex ISH was performed using the manual RNAscope assay (Advanced Cell Diagnostics, ACD). Target-specific probes were obtained from the ACD online catalog: *Slc17a6* (Ref. #319171), *tdTomato* (Ref. #317041), and *Slc32a1* (Ref. #319191). Frozen brain sections were fixed in 4% paraformaldehyde (PFA; Ted Pella, Inc., Cat. #18505) for 15 minutes at room temperature, followed by sequential dehydration in graded ethanol (Hayman, Cat. #64-17-5) for 20 minutes. After brief air drying, a hydrophobic barrier was drawn around each tissue section. RNAscope Hydrogen Peroxide (Cat. #322335) was applied for 10 minutes, followed by two rinses in nuclease-free water (MP Biomedicals, Cat. #112450204). Sections were then incubated with Protease IV (Cat. #322336) for 30 minutes and rinsed twice in nuclease-free water. A mixture of probes was prepared at a ratio of 50:1:1 for *Slc17a6* (channel 1), *tdTomato* (channel 2), and *Slc32a1* (channel 3), and applied to the sections. Hybridization was carried out for 2.5 hours at 40 °C using the HyperChrome hybridization system (HyperChrome, Cat. #EHP 500AS). Following hybridization, sections were washed with 1X wash buffer (Cat. #320058), and signal amplification and chromogenic development were performed according to the manufacturer’s protocol (ACDBIO). Images for anatomical analysis were acquired using 10X and 20X objectives on a Leica SP8 Falcon laser scanning confocal microscope (Leica Microsystems, Germany) and processed using Leica image analysis suite software.

#### Input-output mapping of LHA^stress-TRAP^ neurons

To map the brain-wide monosynaptic inputs of the LHA^stress-TRAP^ neurons, we used the pseudorabies virus-based retrograde tracing strategy^45,46^. Briefly, rAAV5-EF1α-DIO-oRVG and rAAV5-EF1α-DIO-EGFP-T2A-TVA were injected first. Three weeks later, mice were stress-TRAPed as described above, and then a second injection of RV-CS-N2C-deltaG-tdTomato was done one week after the stressTRAPing. Brain tissue was harvested after 1 week of rabies injection for histochemical analysis as described above. Every 3rd brain section was mounted and imaged on the upright fluorescence microscope (Khush Enterprises, Bengaluru) (2X, 4X, and 10X lenses). ImageJ/FIJI processing software was used to process the images, and the number tdTomato positive cells were counted reported from 3 mice.

To map the brain-wide downstream targets of the LHA^stress-TRAP^ neurons, AAV-DIO-tdTomato was injected into the LHA of the TRAP2 mice, and stress-TRAPing was done 3 weeks after the injection. One week after the stress-TRAPing, the brain tissue was harvested, and the brain-wide projections of the LHA^stress-TRAP^ neurons were imaged under the fluorescent microscopy. Gain and exposure time were kept constant throughout the imaging session. A box area was selected in the region containing the tdTomato positive fibers to calculate the density of the projection, and the mean intensity (F_Total_) was calculated using the ImageJ/FIJI. To calculate the background intensity (F_Background_), the same box area was dragged to a region, where tdTomato positive fibers were absent on the same brain section and then the mean intensity was calculated. The mean fluorescent intensity of the region of interest (F_ROI_) was reported as F_ROI_= F_Total_ - F_Background_.

#### Quantification and statistical analysis

All statistical analyses were performed using GraphPad PRISM 8.0.2 software. Student t-test and Two-way ANOVA tests were performed wherever applicable. ns > 0.05, ∗ P ≤ 0.05, ∗∗ P ≤ 0.01, ∗∗∗ P ≤ 0.001, ∗∗∗∗ P ≤ 0.0005.

## Results

### RS suppresses non-histaminergic acute and chronic psoriatic itch

To elucidate the neural circuits for stress-modulation of itch, first, we probed whether acute restraint stress (RS) affects acute and chronic itch in mice. RS is a commonly used method to induce stress and anxiety in rodents and to study stress’s physiological and behavioral effects^4,47–49^. We induced RS by restricting wild-type CD-1 mice in a tube for an hour^4,47^ and injected chloroquine, a non-histaminergic pruritogen, in the nape of the neck (Figure 1A) ^50^. Compared to the unrestrained control mice, the RS suppressed chloroquine-induced acute itch (Figure 1B). Next, we tested if RS suppresses the spontaneous psoriatic itch caused by repeated application of imiquimod on the nape of the neck in CD-1 mice(Figure 1C). We found that RS alleviated the pathological spontaneous itch observed in psoriatic mice compared to unrestrained animals (Figure 1D). Thus, we found that acute stress suppresses both acute and chronic itch in mice.

**Figure 1:**
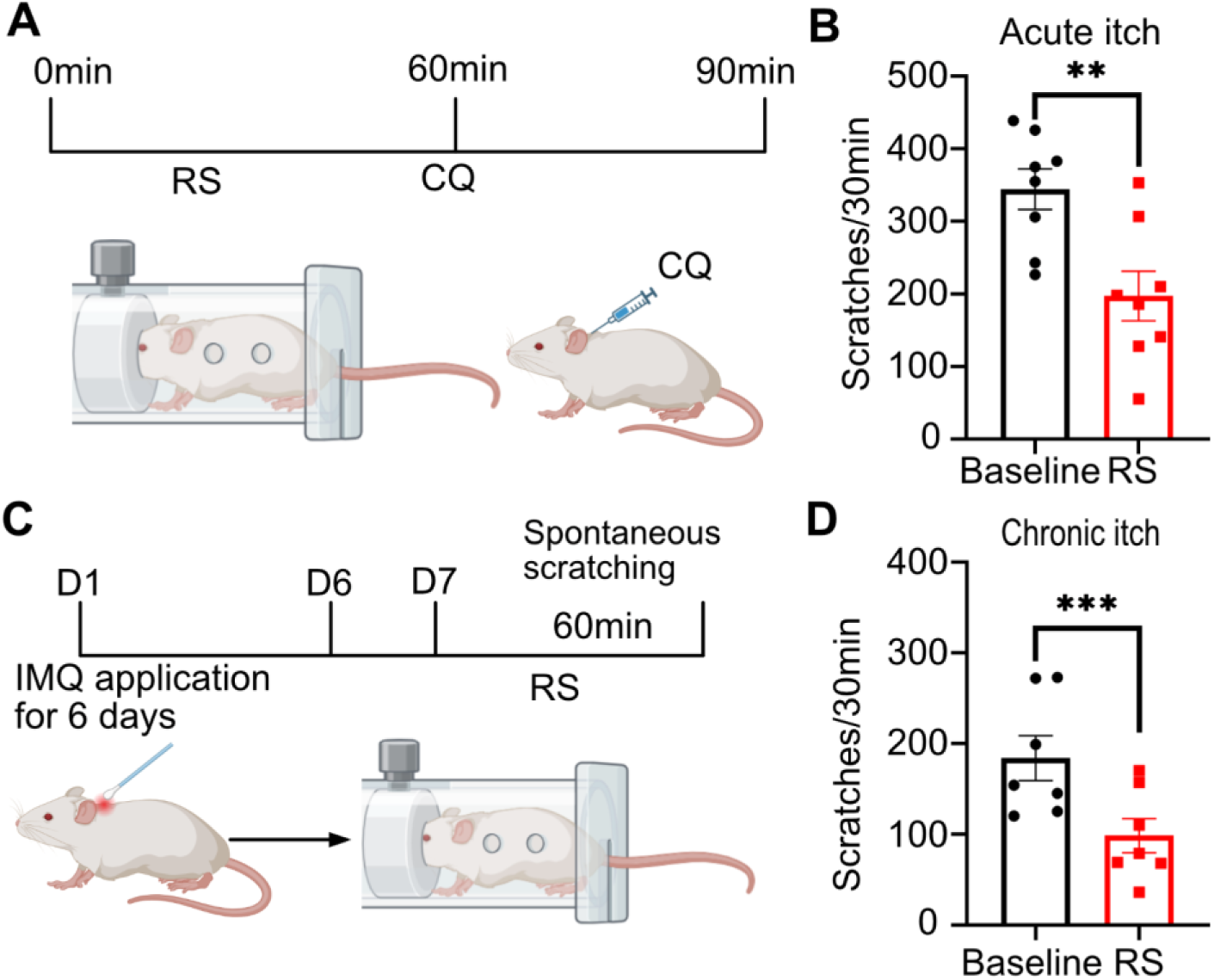
RS suppresses non-histaminergic acute and chronic psoriatic itch. (A) Schematic of the experiment timeline. Mice were restrained in the restraining tube for an hour. Intradermal chloroquine (375 µg / 75 µl) was administered, and scratching behaviour was recorded for 30 minutes. (B) Effect of restraint stress (RS) (t-test, 147 ± 41.31 **P = 0.0092, n = 8) on chloroquine-induced scratching compared with baseline scratching. (C) Schematic of the experiment timeline. Imiquimod (IMQ) was topically applied on the nape of the neck of the mice to induce the psoriatic itch in mice. Mice were restrained for an hour, and then the spontaneous scratching was recorded for 30 minutes. (D) Effect of restraint stress (RS) (t-test, 85.57 ± 12.20 ***P = 0.0004, n = 7) on IMQ-induced spontaneous scratching compared with baseline spontaneous scratching.

### LHA^stress-TRAP^ neurons are sufficient for anxiety-like behaviors in mice

Next, we explored whether we can label stress-sensitive neurons in the LHA with the TRAP2 transgenic strain (Figure S1). We reasoned that the ability to carry out permanent genetic labeling of stress-sensitive neurons with Cre-recombinase in the LHA would enable transient manipulation of neural activity, anatomical mapping of pre- and post-synaptic inputs, and targeted calcium imaging. RS stress, followed by i.p. 4-OHT administration in TRAP2 mice stereotaxically injected with AAV-DIO-tdTomato in the LHA (Figure S1A), labeled a subpopulation of neurons with tdTomato fluorescence (Figure S1B). Re-introducing the LHA^stress-TRAP- tdTomato^ mice to RS induced cFos expression, an IEG, and a proxy for neural activity in the tdTomato-expressing neurons, indicating the efficiency and the specificity in stress-TRAPping of the LHA neurons (Figure S1C). Furthermore, multiplexed in-situ hybridizations (RNAscope) revealed that a large population of labeled neurons expressed the gene encoding the vesicular glutamate transporter, VGlut2 (*slc17a6*), suggesting that LHA^stress-TRAP^ neurons are mostly excitatory and glutamatergic (Figure S2B and C). Next, tested if the transient activation of LHA^stress-TRAP^ neurons would cause behavioral changes in mice. Specifically, we expected that stimulating LHA^stress-TRAP^ neurons would cause anxiety-like behaviors, one of the common behavioral outcomes of the RS assay. To transiently activate the LHA^stress-TRAP^ neurons, we expressed hM3Dq-mCherry, the depolarizing chemogenetic actuator, in a Cre-dependent manner (LHA^stress-TRAP-hM3Dq^) (Figure 2A). I.p. administration of deschloroclozapine (DCZ) activated the LHA^stress-TRAP^ neurons, and was reflected in the expression of cFos in the LHA^stress-TRAP-hM3Dq^ neurons expressing mCherry (fused to hM3Dq, which helps to visualize hM3Dq expression) (Figure 2B). We found that DCZ in LHA^stress-TRAP-hM3Dq^ mice and not in control mice expressing tdTomato in the LHA (LHA^stress-TRAP-tdTomato^) promoted anxiety-like behaviors as assayed on the open-field test (OFT), where mice with anxiety tend to spend less time in the center zone, and on the light-dark box (LDB) test, where anxious mice preferred to stay in the dark box. Together, our behavioral data indicate that the chemogenetic activation of the LHA^stress-TRAP^ neurons is sufficient to promote stress and anxiety (Figure 2C-K).

**Figure 2:**
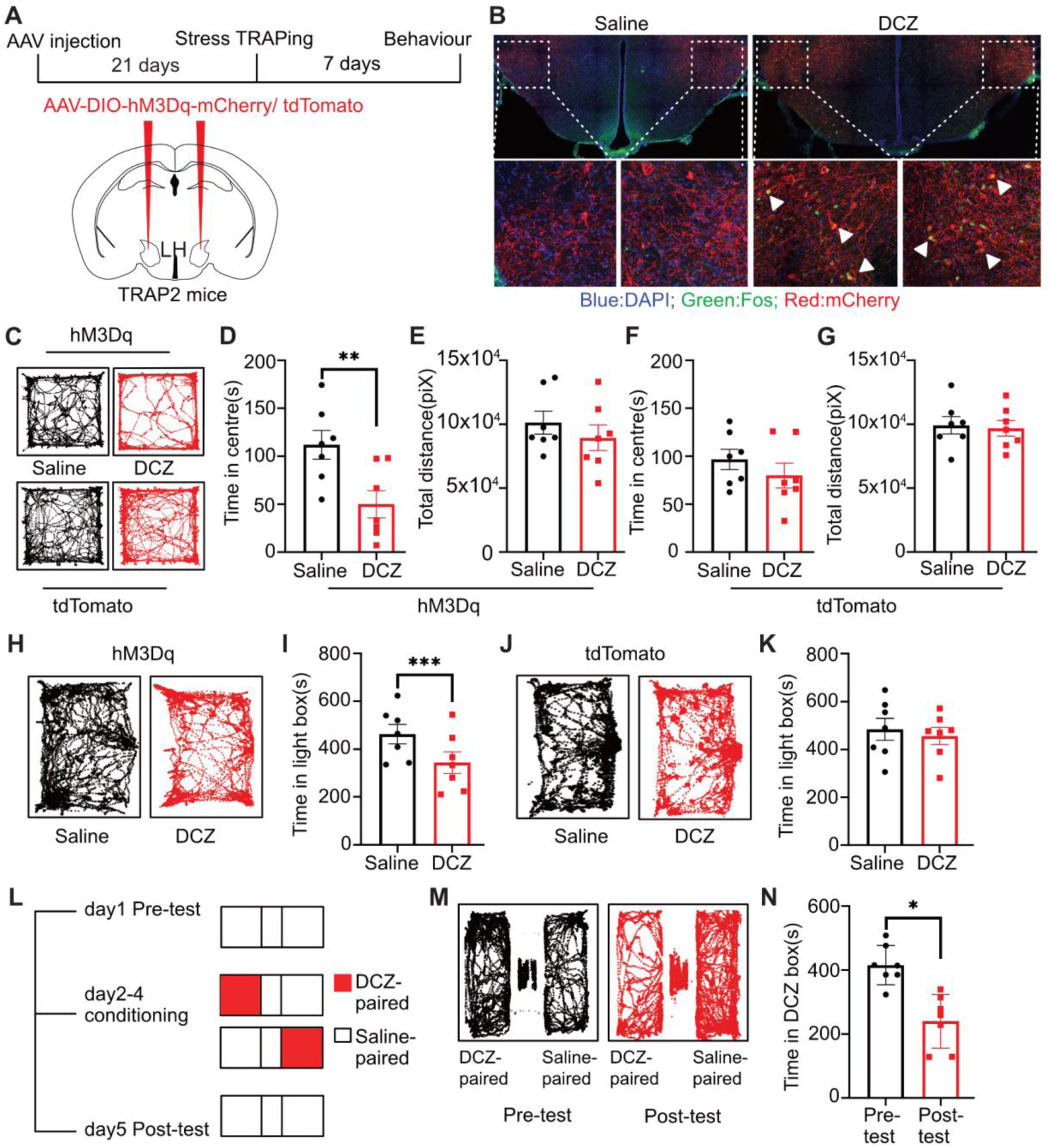
LHA^stress-TRAP^ neurons are sufficient for anxiety-like behaviors in mice. (A) Schematic of the experiment timeline. AAV encoding Cre-dependent excitatory DREADD hM3Dq or tdTomato was injected bilaterally into the LHA of the TRAP2 mice. (B) Coronal section image of LHA from the injected TRAP2 mice shows the expression of hM3Dq (red: mCherry) in LHA^stress-TRAP^ neurons. DCZ administration results in Fos(green) induction in the hM3Dq expressing neurons. White triangles show yellow cells expressing both mCheery and Fos. (C) Example trajectories of mice in the open field arena after saline/DCZ administration. (D) Effect of DCZ on time spent in the centre zone (t-test, 62.07 ± 10.52 **P = 0.0011, n = 7) compared to saline administration in the hM3Dq injected mice. (E) The total distance travelled in the arena was unaffected after DCZ administration as compared to saline administration in the hM3Dq injected mice. (F) Time spent in the centre zone was unaffected between DCZ and saline administered conditions in the tdTomato injected mice. (G) The total distance travelled in the arena was unaffected between DCZ and saline administered conditions in the tdTomato injected mice. (H) Example trajectory of a mouse in the light box after saline/DCZ administration in the light-dark box test. (I) Effect of DCZ on time spent in light box (t-test, 118.8 ± 13.49 ***P = 0.0001, n = 7) compared to saline administration in the hM3Dq injected mice. (J) Example trajectory of a mouse in the light box after saline/DCZ administration in the light-dark box test in the tdTomato injected mouse. (K) Time spent in the light box after DCZ administration was unaffected as compared to saline administration in the tdTomato injected mice. (L) Experimental strategy for the conditioned place aversion (CPA) test. (M) Example trajectory of a mouse pre- and post-conditioning in the CPA apparatus. (N) Effect of DCZ on time spent in DCZ paired chamber (t-test, 176 ± 48.05 *P = 0.0105, n = 7) compared to saline administration.

Further, we determined if transient activation of the LHA^stress-TRAP^ neurons is enough to cause learned aversion, which was tested through the conditioned place aversion (CPA) test. In the CPA test, where the conditioning stimulus is the chemogenetic activation of a neuronal population of interest, the experimental animals are paired with a chamber with i.p. DCZ and asked if the pairing is sufficient to drive aversion. We conditioned one of the chambers of the CPA apparatus with DCZ in a cohort of LHA^stress-TRAP-hM3Dq^ mice and the other chamber with i.p. saline (Figure 2L). We found that the LHA^stress-TRAP-hM3Dq^ mice spent less time in the DCZ-paired chamber on the test day (Figure 2M and N). Thus, activation of LHA^stress-TRAP^ neurons are sufficient to drive CPA.

### LHA^stress-TRAP^ neurons bidirectionally control acute and chronic itch

In the next set of experiments, we tested the effect of chemogenetic activation of the LHA^stress-TRAP^ neurons on acute and chronic itch (Figure 3A). As mentioned before, experimental acute itch was induced by intradermal injection of chloroquine (Figure 1A and B). In contrast, chronic itch was modeled by repeated application of imiquimod on the nape of the neck of mice (Figure 1C, D). We found that i.p. DCZ-mediated stimulation of the LHA^stress-TRAP-hM3Dq^ neurons suppressed both acute and chronic itch (Figure 3B and C), while the same manipulation did not affect acute and chronic itch in the tdTomato injected control mice (Figure S3G and H). We previously showed that the LHA neurons downstream of the lateral septum (LS) mediate SIA. Hence, we wondered if activating the LHA^stress-TRAP^ neurons enhanced spinal reflexive and supra-spinal thermal pain thresholds. We used the tail-flick test to determine the spinal reflexive thermal nociceptive thresholds and to assay the supraspinal thresholds and nocifensive behaviors; we used the hot-plate assay. We found that transient activation of the LHA^stress-TRAP^ neurons increased spinal and supraspinal thermal thresholds (Figure S3A-E), while nociceptive thresholds remained unchanged in the control mice (Figure S3I-M). Thus, LHA^stress-TRAP^ neurons mediate stress-mediated suppression of pain and itch. Next, we wondered how silencing the LHA^stress-TRAP^ neurons would affect acute and chronic itch. To that end, we expressed the inwardly rectifying potassium Kir2.1 channel in the LHA^stress-TRAP^ neurons in a Cre-dependent manner (LHA^stress-TRAP-Kir2.1^)(Figure 3D and E). We injected AAV-DIO-GFP in the LHA of stress-TRAP mice and used them as controls. Silencing the LHA^stress-TRAP^ neurons increased acute itch induced by chloroquine and imiquimod-induced psoriasis compared to control mice expressing GFP (Figure 3F and G). Thus, the LHA^stress-TRAP^ neurons are necessary for maintaining normal scratching frequency in response to pruritogens and chronic inflammatory itch. We then asked if the LHA^stress-TRAP^ neurons are required for stress-induced suppression of acute and chronic itch. We found that the Kir2.1-mediated silencing of the LHA^stress-TRAP^ neurons reduced the suppressive effects of RS on acute and psoriatic itch (Figure 3J and L). At the same time, in GFP-expressing control mice, RS inhibited itch, as seen previously (Figure 3I and K). In conclusion, the LHA^stress-TRAP^ neurons are required for acute stress-mediated itch suppression.

**Figure 3:**
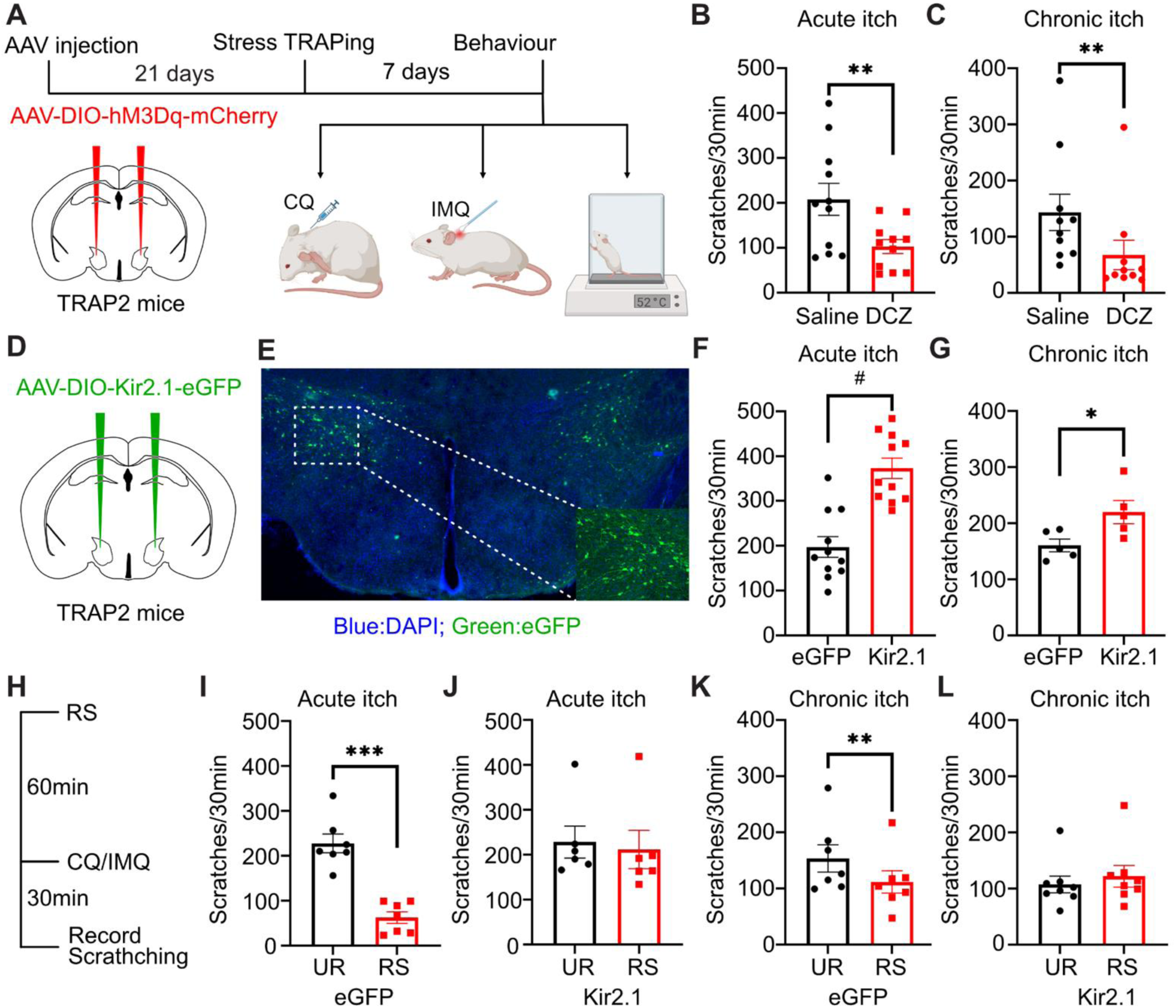
LHA^stress-TRAP^ neuron activation suppresses acute and chronic itch. (A) Schematic of the experiment timeline. AAV encoding Cre-dependent excitatory DREADD hM3Dq was injected bilaterally into the LHA of the TRAP2 mice. (B) Effect of i.p. DCZ (t-test, 105.2 ± 31.43, **P = 0.0074, n = 11) administration on chloroquine-induced scratching compared with i.p. saline. (C) Effect of i.p. DCZ (t-test, 76 ± 19.99, **P = 0.0042, n = 10) administration on psoriatic itch-induced spontaneous scratching compared with i.p. saline. (D) AAV encoding Cre-dependent Kir2.1 was injected bilaterally into the LHA of the TRAP2 mice and stress-TRAPed after 21 days. (E) Coronal section of LHA showing the expression of Kir2.1 marked by the expression of eGFP (green, DAPI-blue) in the LHA^stress-TRAP^ neurons. (F) Effect of Kir2.1 silencing of LHA^stress-TRAP^ neurons (unpaired t-test, 175.5 ± 32.49, #P = 0.0001) on chloroquine-induced scratching compared with eGFP controls. (G) Effect of Kir2.1 silencing of LHA^stress-TRAP^ neurons (unpaired t-test, 59.20 ± 20.05, *P = 0.0184) on psoriatic itch-induced spontaneous scratching compared with eGFP controls. (H) Schematic of the experiment timeline for testing the necessity of LHA^stress-TRAP^ neurons in stress-induced suppression of itch. (I) Effect of restraint stress (RS) (t-test, 165.4 ± 18.31 ***P = 0.0001, n = 7) on chloroquine-induced scratching compared with unrestrained(UR). (J) Effect of restraint stress (RS) on chloroquine-induced scratching compared with unrestrained(UR) in Kir2.1 mediated silenced LHA^stress-TRAP^ neurons mice. (K) Effect of restraint stress (RS) (t-test, 42 ± 9.86 **P = 0.0053, n = 7) on psoriatic itch-induced spontaneous scratching compared with unrestrained(UR). (L) Effect of restraint stress (RS) on psoriatic itch-induced spontaneous scratching compared with unrestrained(UR) in Kir2.1 mediated silenced LHA^stress-TRAP^ neurons mice.

### Input-output mapping of LHA^stress-TRAP^ neurons

The advent of virally mediated anatomic tracing techniques has revolutionized the mapping of inputs and outputs of select neurons in the central nervous system. Hence, the monosynaptic rabies tracing technique was used here to delineate the presynaptic inputs in the brain. For the monosynaptic rabies tracing, we stereotaxically delivered the Cre-dependent AAVs carrying the genes for the TVA-GFP and RVΔG in the LHA of the TRAP2 mice. After three weeks, we trapped the stress-sensitive neurons in the LHA with 4-OHT to enable TVA-GFP and RVdelG expression in the LHA^stress-TRAP^ neurons. One week later, we injected the RV-N2C-delG-nlstdTomato in the LHA (Figure 4A). We found that in a week, LHA^stress-TRAP^ neurons express the TVA-tagged GFP (green) and the tdTomato (red) from modified rabies viruse, indicating that the double-fluorescently labeled yellow cells are the starter cells (Figure 4C). We observed neurons with red fluorescence in the pruriceptive brain areas such as the LPBN, PAG, and RVM; somatosensory information processing areas such as the deeper layers of the S1; and affective-motivational brain nuclei such as the LC, BLA, BNST, LS, and MS (Figure 4D and E). Previously, we found that the inhibitory neurons in the LS project to the LHA to drive RS-induced analgesia^4^. Since we found that the LS and the LHA^stress-TRAP^ neurons are monosynaptically connected, we tested whether the LHA^stress-TRAP^ neurons are the same as the LHA_post-LS_ cells. To that end, we injected the anterograde transsynaptic AAVTranssyn-FlpO in the LS of TRAP2 mice, facilitating the availability of the FlpO recombinase specifically in the LHA_post-LS_ neurons. Simultaneously, we delivered DIO-GFP and fDIO-tdTomato in the LHA (Figure S4A). After stress trapping, the LHA_post-LS_ and the LHA^stress-TRAP^ neurons expressed tdTomato and GFP, respectively. We found that a few of the GFP and tdTomato-expressing neurons overlap (Figure S4B and C). This implies that the majority of the LHA^stress-TRAP^ neurons do not receive input from the LS. This is in line with our previous findings, where the inhibitory neurons were engaged by RS and suppressed LHA activity^4^. Together, unbiased retrograde mapping of monosynaptic inputs implies that the negative-affective signals due to RS might be routed through the fore and mid-brain limbic areas or brainstem nuclei encoding aversive sensory stimuli.

**Figure 4:**
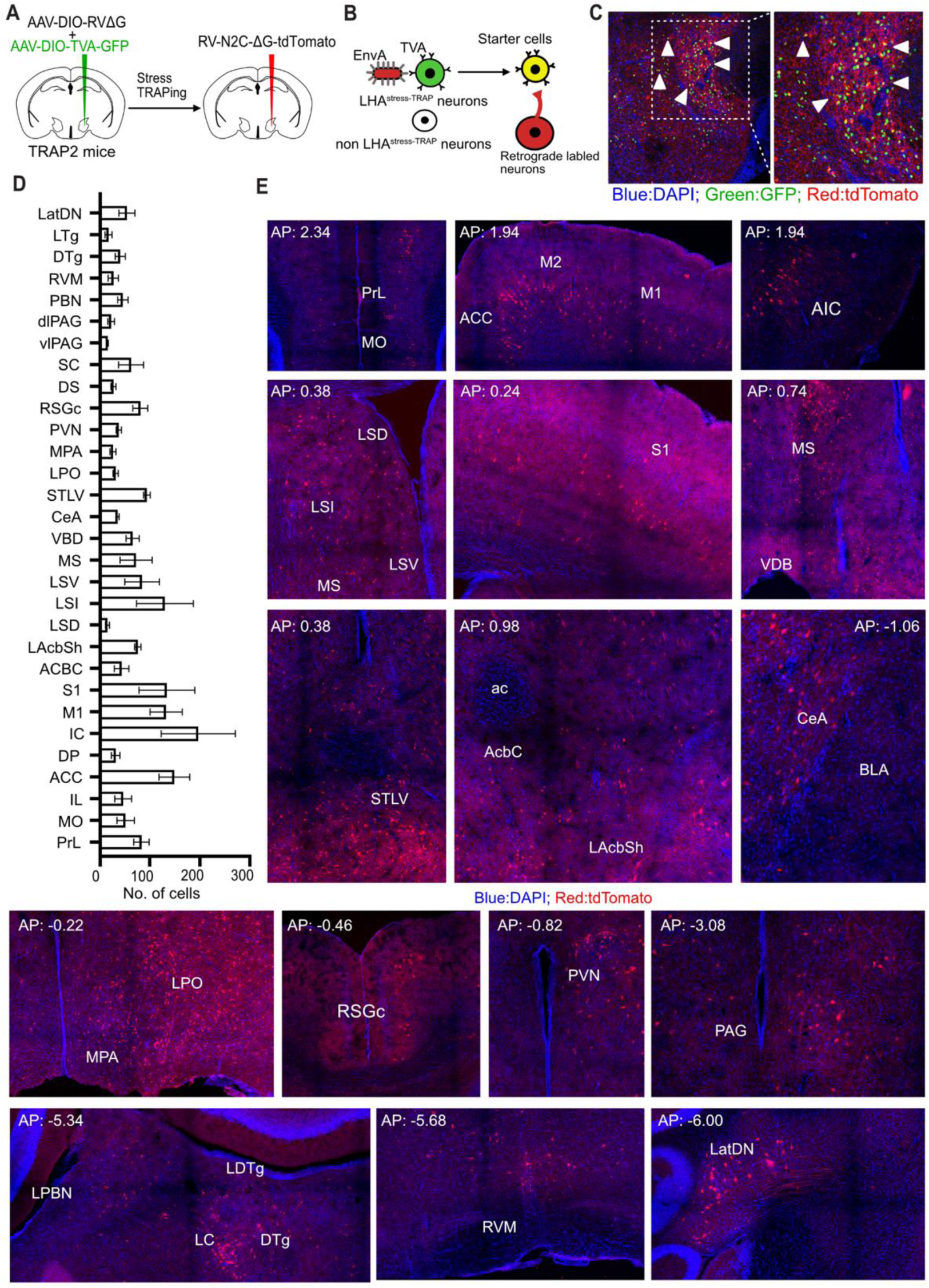
Monosynaptic rabies tracing of the LHA^stress-TRAP^ neurons. (A) Viral strategy to map monosynaptic inputs of the LHA^stress-TRAP^ neurons. Two AAVS-one encoding Cre-dependent rabies glycoprotein G and another encoding avian TVA receptor were mixed into 1:1 ratio and injected into the LHA of the TRAP2 mice. One week after stress TRAPing modified rabies virus with deleted glycoprotein G was injected into the LHA of the same mice. (B) Schematic for the monosynaptic inputs of the LHA^stress-TRAP^ neuron. (C) Confocal image of the LHA showing the starter cells (yellow, marked by white triangles). (D) Whole brain quantification of the inputs to the LHA^stress-TRAP^ neurons. Data are presented as mean ± SEM, from n=3 mice. (E) Representative images of retrogradely–labeled neurons in different brain regions.

Next, we sought to map the brain-wide axonal projections of the LHA^stress-TRAP^ neurons. To that end, we injected the cell-filling AAV-DIO-tdTomato in the LHA of the TRAP2 mice and stress-TRAPed after 3 weeks (Figure 5A). We observed robust expression of tdTomato in the LHA^stress-TRAP^ neurons (Figure 5B). Whole-brain sectioning and confocal imaging revealed LHA^stress-TRAP^ projections in diverse brain areas, including the IL, various subnuclei of the septum, bed nucleus of stria terminalis, central amygdala, paraventricular thalamus, and lateral habenula. In the brainstem, it projects to PAG, VTA, PBN, RVM, and reticular formation (Figure 5C and D). Intriguingly, through anterograde and retrograde tracing experiments, we realised that the LHA^stress-TRAP^ neurons have reciprocal connections with brain nuclei such as the central amygdala, LPBN, and RVM.

**Figure 5:**
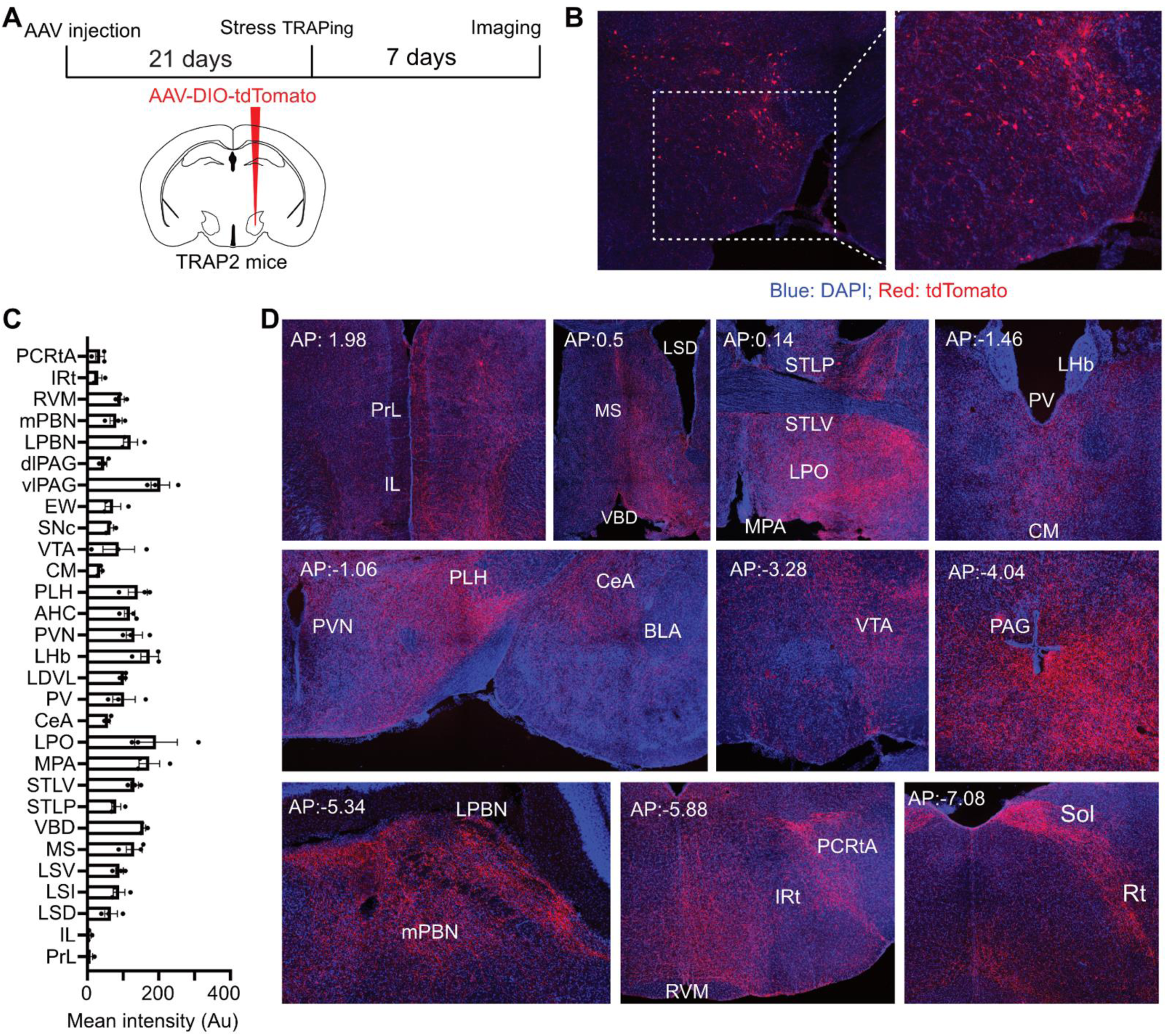
Projections of the LHA^stress-TRAP^ neurons. (A) Schematic of the experiment timeline. AAV encoding Cre-dependent tdTomato was injected bilaterally into the LHA of the TRAP2 mice. One week after stress-TRAPing, brain tissue was harvested for imaging. (B) Confocal image of the injection site. LHA^stress-TRAP^ neurons express tdTomato (red; DAPI-blue). A zoomed-in image of the cells is shown on the right. (C) Whole brain quantification of projection intensity of the LHA^stress-TRAP^ neurons. Data are presented as mean ± SEM, from n=3 mice. (D) Projections of LHA^stress-TRAP^ neurons across different brain regions.

### The activity of LHA^stress-TRAP^ neurons coincides with spontaneous scratching in psoriatic conditions

Genetically encoded fluorescent calcium sensors, such as GCaMP6s, and in vivo calcium imaging techniques, such as fiber photometry, have revolutionized in vivo monitoring of neural activity in behaving mice. Here, we tested the activity of LHA^stress-TRAP^ neurons when the mice were under RS. We injected the AAV9-DIO-GCaMP8s in the LHA of the TRAP2 mice and expressed the GCaMP8s in the acute stress-sensitive neurons by i.p. administration of 4-OHT and then exposing the mice to the RS assay (Figure 6A). Next, we implanted a 200 μm inner diameter fiber optic cannula in the LHA so that the fluorescent dynamics of the GCaMP8s sensor expressed in the LHA^stress-TRAP^ neurons can be recorded with a fiber photometry setup (Figures 6A right panel). Predictably, we found that the LHA^stress-TRAP^ neurons were engaged when the mice were restrained, and specifically, the rise in GCaMP8s fluorescence levels coincided with the bouts of struggle (Figures 6C and D). Similarly, when mice were hung by their tails, a commonly used mild stress stimulus, the LHA^stress-TRAP^ neurons were active (Figure 6E and F). However, in spite of the inhibitory effects of the LHA^stress-TRAP^ neurons on acute and chronic itch (Figure 3), the activity of these neurons did not coincide with scratching bouts induced by chloroquine (Figure 6G, H, and M). Similarly, the activity of the LHA^stress-TRAP^ neurons did not coincide with the nocifensive behaviors, such as licks and shakes on the 52 °C thermal-plate test (Figure S5 A-F). The thermal-plate test enables testing of rodent behavior exposed to a range of surface temperatures, hot and cold, at both innocuous and noxious ranges. Exposed to unbearable, noxious heat above 44 °C, mice and rats respond with reflexive shaking and coping licking responses. Thus, the LHA^stress-TRAP^ neurons are tuned to the stimuli with the potential to cause stress and anxiety; however, they are not activated while mice experience and react to aversive and noxious somatosensory stimuli, causing itch and pain. Next, we tested if the activity of the LHA^stress-TRAP^ neurons is altered in psoriatic conditions and if the pathological spontaneous scratches correlate with the neural activity (Figure 6I, J, and N). To our surprise, we found that neural activity in the LHA^stress-TRAP^ cells correlated with spontaneous (alloknesis) and chloroquine-evoked (hyperkinesis)^51^ in mice with imiquimod-induced psoriasis (Figure 6K, L, and O). This finding is in contrast to our observation that the activity of the LHA^stress-TRAP^ neurons does not coincide with chloroquine-induced acute scratching (Figure 6G and H). Thus, through fiber-photometry recordings, we found that the LHA^stress-TRAP^ neurons are activated by stressors and engaged by itch under chronic psoriatic conditions.

**Figure 6:**
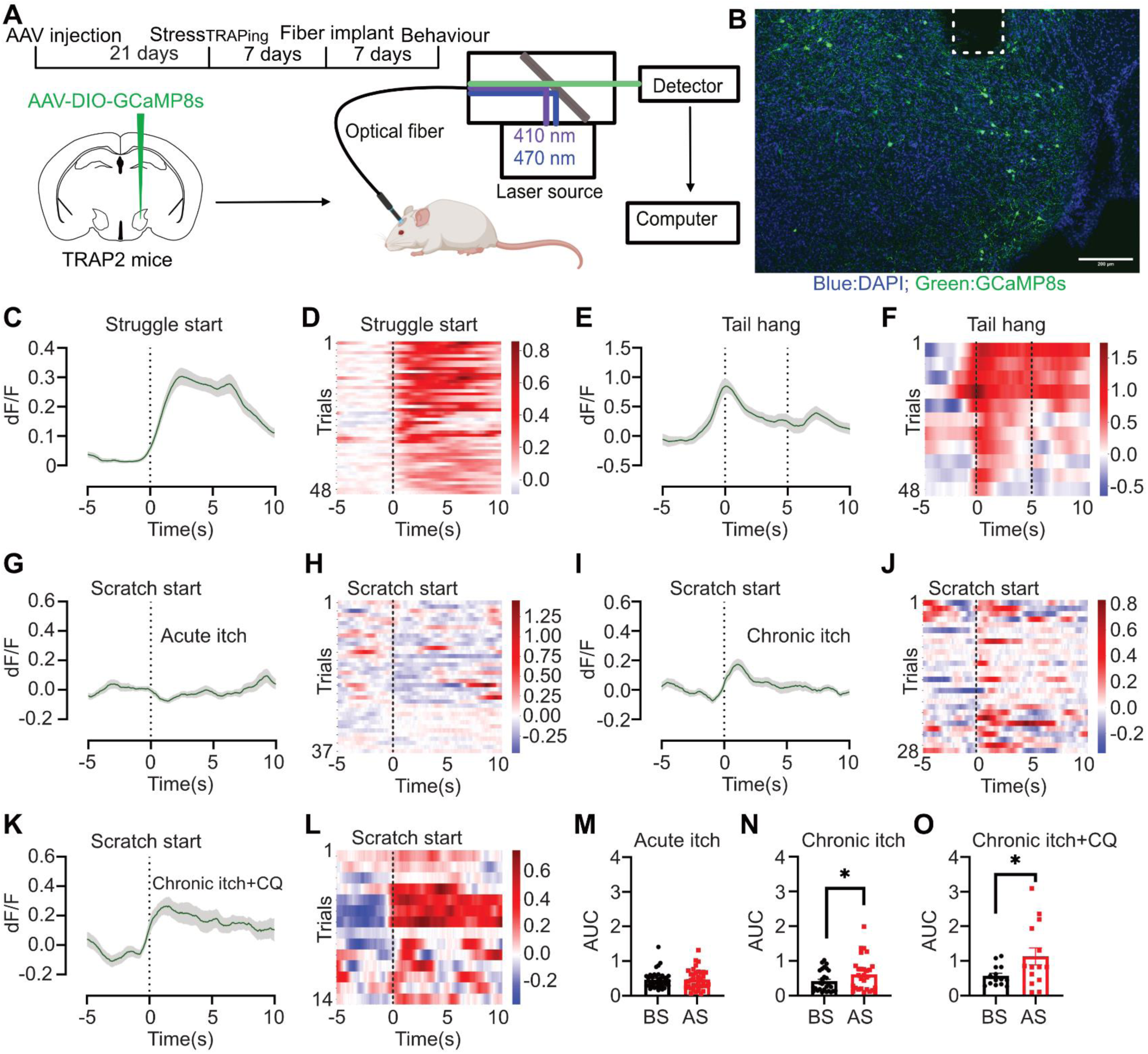
LHA^stress-TRAP^ neurons respond to chronic itch-induced scratching. (A) Schematic of the experiment. AAV encoding Cre-dependent GCaMP8s was injected into the LHA of the TRAP2 mice. The right panel shows the schematics of dual colour photometry. (B) Coronal section image of LHA from the injected TRAP2 mice shows the expression of GCaMP8s (green; DAPI: blue) in LHA^stress-TRAP^ neurons. The white dotted line shows the fiber track. (C) The average fluorescent signal at the start of the struggle in the restraint stress. (D) Heatmap showing the response to struggle in the restraint stress. (E) The average fluorescent signal during the tail hang test. (F) Heatmap showing the response during the tail hang test. (G) The average fluorescent signal at the start of chloroquine-induced scratching. (H) Heatmap showing the response at the start of chloroquine-induced scratching. (I) The average fluorescent signal at the start of spontaneous scratching in psoriatic chronic itch. (J) Heatmap showing the response of spontaneous scratching in psoriatic chronic itch. (K) The average fluorescent signal at the start of scratching induced by chloroquine in the psoriatic chronic itch. (L) Heatmap showing the response of scratching induced by chloroquine in the psoriatic chronic itch. (M) Area under the curve (AUC) 5s before and after the start of chloroquine induced scratching. (N) Area under the curve (AUC) 5s before and after the start spontaneous scratching in chronic itch (t-test, 0.1993 ± 0.08799, *P = 0.0318, n = 28). (O) Area under the curve (AUC) 5s before and after the start of chloroquine induced scratching in chronic itch condition (t-test, 0.5633 ± 0.2047, *P = 0.0165, n = 14).

### LHA^stress-TRAP^ neurons are potentiated in psoriatic mice

In vivo recordings revealed that the LHA^stress-TRAP^ neurons become responsive to pruritic stimuli in mice that develop experimental psoriasis (Figure 6K, L, and O). Thus, we hypothesized that imiquimod may sensitize the LHA^stress-TRAP^ neurons and that should be reflected in increased firing rates in ex vivo preparations with postsynaptic current injections. To test, we performed whole-cell patch-clamp electrophysiological recordings in fluorescently labelled (eGFP) LHA^stress-TRAP^ neurons from mice with and without psoriasis (controls) (Figure 7A and B). We expressed eGFP in the LHA^stress-TRAP^ neurons using the method described in previous sections (Figure S1). Indeed, we observed a gain in firing in LHA neurons of imiquimod-induced psoriasis mice compared to neurons in control mice that is reflected in the F/I (frequency/current) plot by an increase in the frequency of action potentials upon a series of current injections (Figure 7D) ranging from 0 pA to 300 pA postsynaptic depolarizing current injections with 20pA increments. Similarly, we observed a significant hyperpolarizing shift in the rheobase membrane voltage, implying higher excitability propensity of LHA^stress-TRAP^ neurons of psoriasis mice compared to the controls (Figure 7E). Furthermore the latency to first spike firing had decreased in the psoriasis mice compared to the controls (Figure 7F). Meanwhile, the action potential amplitude, rise, and decay time constants remained unaffected between psoriasis and control neurons (Figure 7G-I). Notably, the passive membrane properties, like input resistance and the membrane voltage, didn’t change between the psoriasis and control mice (Figure S6A and B), indicating an intrinsic change in membrane excitability contributing to higher excitability of LHA^stress-TRAP^ neurons in psoriasis mice.

**Figure 7:**
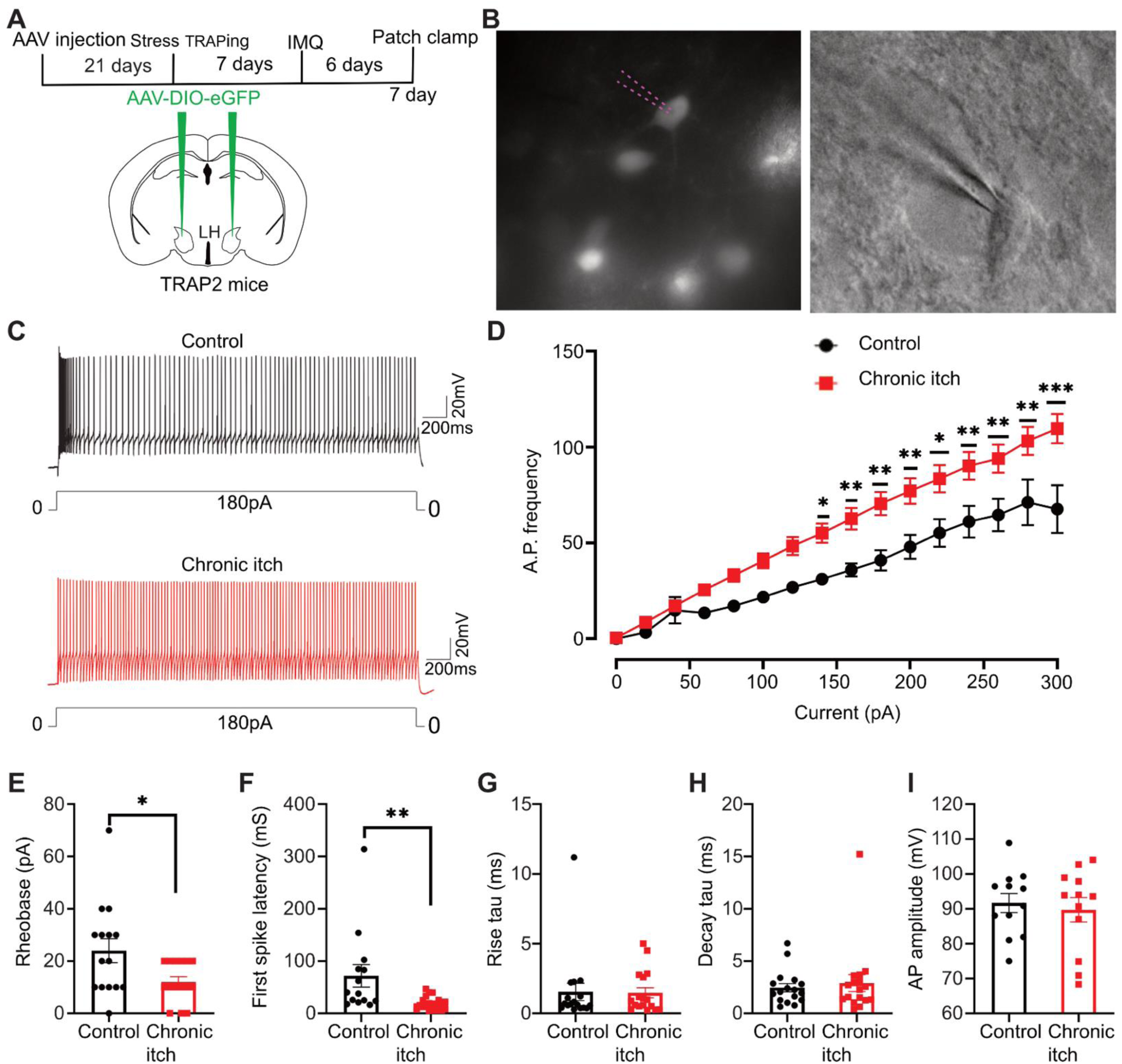
LHA^stress-TRAP^ neurons become hyperactive after chronic itch. (A) Schematic of the experiment timeline. AAV encoding Cre-dependent tdTomato was injected bilaterally into the LHA of the TRAP2 mice. (B) Image of a representative patched cell. Dotted lines show the pipet tip. (C) Representative trace of AP firing in control and chronic itch mice, from the LHA^stress-TRAP^ neurons. (D) Effect of chronic itch on the frequency of AP as compared to control AP mice. Two-way ANOVA (n = 17 cell for chronic itch from 6 mice and n = 12 cells from 5 control mice, P values from 140pA to 300pA currents are 0.0274, 0.008,.0071,0.0053, 0.0122, 0.0086, 0.0081, 0.0054, 0.0001). (E) Effect of chronic itch on rheobase (unpaired t-test, 12 ± 4.976, *P = 0.0227) of the LHA^stress-TRAP^ neurons as compared to the controls. (F) Effect of chronic itch on first spike latency of the LHA^stress-TRAP^ neurons (unpaired t-test, 52.84 ± 18.66, *P = 0.0081) (G) Chronic itch had no effect on the rise tau of the LHA^stress-TRAP^ neurons as compared to the controls. (H) Chronic itch had no effect on the decay tau of the LHA^stress-TRAP^ neurons as compared to the controls. (I) Chronic itch had no effect on the action potential amplitude of the LHA^stress-TRAP^ neurons as compared to the controls.

### LHA^stress-TRAP^ neurons suppress acute and chronic itch through axonal projections in the PAG and RVM

LHA^stress-TRAP^ neurons were sufficient to suppress acute and chronic psoriatic itch (Figure 3). Hence, we sought to understand the downstream target brain nuclei through which LHA^stress-TRAP^ neurons mediate the expression of nocifensive scratching responses to pruritic stimuli. Anterograde tracing revealed that LHA^stress-TRAP^ neurons project to regions such as PAG, PBN, and RVM, which are known to modulate itch-induced scratching through direct or indirect projections to the spinal cord (Figure 5). Notably, these brainstem nuclei are interconnected, receive hypothalamic inputs, and are known to determine modulation of nociceptive and itch thresholds by internal brain states such as stress and hunger^4,39,52,53^. We expressed hM3Dq-mCherry in the LHA^stress-TRAP^ neurons (LHA^stress-TRAP-hM3Dq^) using the genetic strategy described before (Figure 2), and implanted cannulae in the axonal target regions such as the PAG, LPBN, and RVM to enable localized DCZ infusion and specific excitation of the downstream neurons. We implanted bilateral cannulae over PAG and LPBN and single cannulae over RVM due to the central location of the nuclei. Successful expression of hM3Dq-mCherry in the LHA^stress-TRAP^ neurons and labeling of axon terminals in the target regions was evidenced by the presence of mCherry (red) terminals. While chemogenetic activation of the PAG, LPBN, and RVM target neurons by DCZ infusion was confirmed by cFos expression (Figure 8B). As controls for the targeted chemogenetic activation of LHA^stress-TRAP^ downstream target neurons, we expressed tdTomato in the LHA^stress-TRAP^ (LHA^stress-TRAP-tdTomato^) neurons. Thus, in the control mice, DCZ infusion in the LHA^stress-TRAP-tdTomato^ mice would not activate neurons in the LHA, PAG, LPBN, or RVM. We found that the DCZ infusion through cannulae in PAG and RVM of the LHA^stress-TRAP-hM3Dq^ mice, and not in the LPBN, suppressed intradermal chloroquine-induced itch in the nape of the neck (Figure 8C, G, and K). Since chemogenetic activation of the LHA^stress-TRAP^ neurons resulted in analgesia on the hot-plate and tail-flick tests, and LHA-RVM circuitry is known to modulate nociceptive thresholds, we tested if DCZ infusion in the RVM of the LHA^stress-TRAP-hM3Dq^ mice would affect nocifensive behaviors to thermal stimuli. We found that transient activation of LHA^stress-TRAP^ terminals in the RVM was sufficient for increasing the latency to lick and shake on the hot-plate test, increasing the frequency of occurrences of both the nocifensive behaviors, as well as resulting in elevated thresholds on the tail-flick test (Figure S6C-G). In LHA^stress-TRAP-tdTomato^ mice, DCZ at PAG, LPBN, or RVM did not alter scratching by chloroquine (Figure 8E, I, and M). At the same time, spontaneous scratching caused by repeated application of imiquimod was suppressed by chemogenetic activation of axon terminals of the LHA^stress-TRAP-hM3Dq^ neurons in the PAG and RVM, and not LPBN (Figure 8D, H, and L). DCZ infusion in either of the downstream targets of the LHA^stress-TRAP^ neurons in the tdTomato expressing mice did not alter the psoriasis-induced spontaneous scratching (Figure 8F, J, and N). Thus, the LHA^stress-TRAP^ neurons, through their downstream targets in the PAG and RVM, mediate the effects of stress on acute and chronic itch.

**Figure 8.**
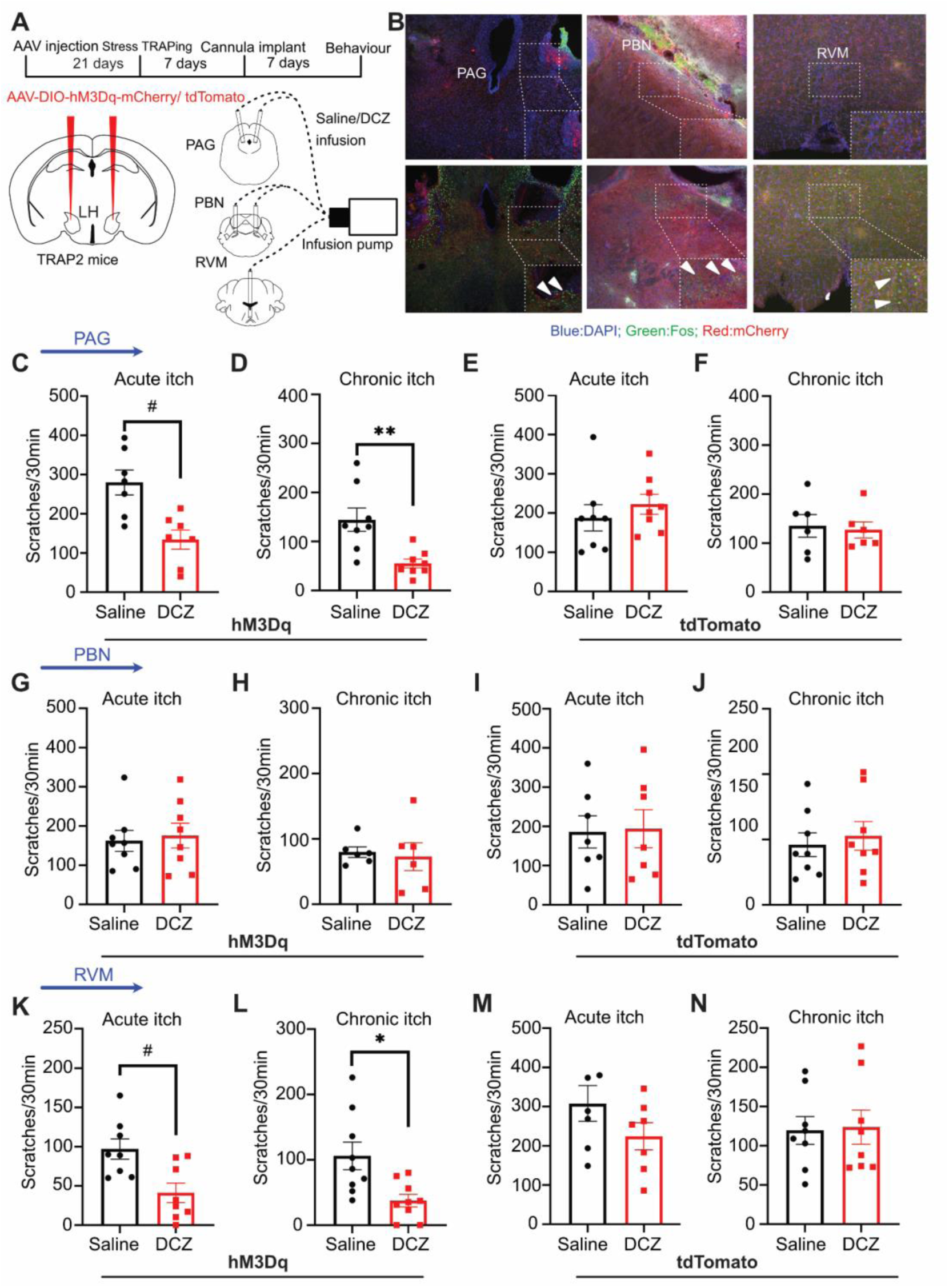
LHA^stress-TRAP^ neurons suppress acute and chronic itch through axonal projections in the PAG and RVM. (A) Schematic of the experiment timeline. AAV encoding Cre-dependent excitatory DREADD hM3Dq was injected bilaterally into the LHA of the TRAP2 mice. One week after stress TRAPing bilateral cannula was implanted at the vlPAG, PBN, and RVM. (B) Coronal section image of PAG, PBN, and RVM. DCZ infusion results in Fos (green) induction in mCherry (red) surrounded PAG, PBN, and the RVM neurons. (C) Effect of DCZ infusion at PAG (t-test, 145.9 ± 13.97, #P < 0.0001, n = 7) on chloroquine-induced scratching compared with saline infusion. (D) Effect of DCZ infusion at PAG (t-test, 89.50 ± 17.02, **P = 0.0012, n = 8) on psoriatic itch induced spontaneous scratching compared with saline infusion. (E) DCZ infusion at PAG has no effect on chloroquine-induced scratching compared with saline infusion in tdTomato injected control mice. (F) DCZ infusion at PAG on psoriatic itch induced spontaneous scratching compared with saline infusion in tdTomato injected control mice. (G) DCZ infusion at PBN has no effect on chloroquine-induced scratching compared with saline infusion. (H) DCZ infusion at PBN has no effect on psoriatic itch-induced spontaneous scratching as compared with saline infusion. (I) DCZ infusion at PBN has no effect on chloroquine-induced scratching compared with saline infusion in tdTomato injected control mice. (J) DCZ infusion at PBN has no effect on psoriatic itch-induced spontaneous scratching as compared with saline infusion in tdTomato injected control mice. (K) Effect of DCZ infusion at RVM (t-test, 55.75 ± 6.761, #P < 0.0001, n = 8) on chloroquine-induced scratching compared with saline infusion. (L) Effect of DCZ infusion at RVM (t-test, 68.56 ± 21.86, *P = 0.0139, n = 9) on psoriatic itch induced spontaneous scratching compared with saline infusion. (M) DCZ infusion at RVM has no effect on chloroquine-induced scratching compared with saline infusion in tdTomato injected control mice. (N) DCZ infusion at RVM has no effect on psoriatic itch-induced spontaneous scratching as compared with saline infusion in tdTomato injected control mice.

Further, we tested if the optogenetic inhibition of the axon terminals of the LHA^stress-TRAP^ neurons in the PAG altered acute and chronic itch. To that end, we expressed the halorhodopsin fused with YFP (eNpHR3.0-YFP)^54^ in a Cre-dependent manner in the LHA^stress-TRAP-eNpHR3.0^ neurons with AAV vectors delivered stereotaxically following the strategy described above (Figure 9A). We expressed YFP in the LHA^stress-TRAP^ neurons and used them as controls. The expression of eNpHR in the LHA^stress-TRAP^ neurons was confirmed by the expression of YFP, which was fused to the eNpHR (Figure 9B). To optogenetically inhibit the LHA^stress-TRAP-eNpHR3.0,^ we implanted bilateral optic fiber cannula in the PAG of mice (Figure 9C). Yellow light (595 nm) shined through the cannulae in PAG exacerbated chloroquine-induced acute and imiquimod-induced chronic itch (Figure 9D and E). Predictably, in control experiments, the yellow light shined at the YFP-expressing terminals of the LHA^stress-TRAP^ neurons in the PAG did not alter acute and chronic itch (Figure 9F and G). In sum, the suppression of the LHA^stress-TRAP^ inputs to the PAG enhances the urge to scratch the site of itch under acute and chronic pathological conditions.

**Figure 9.**
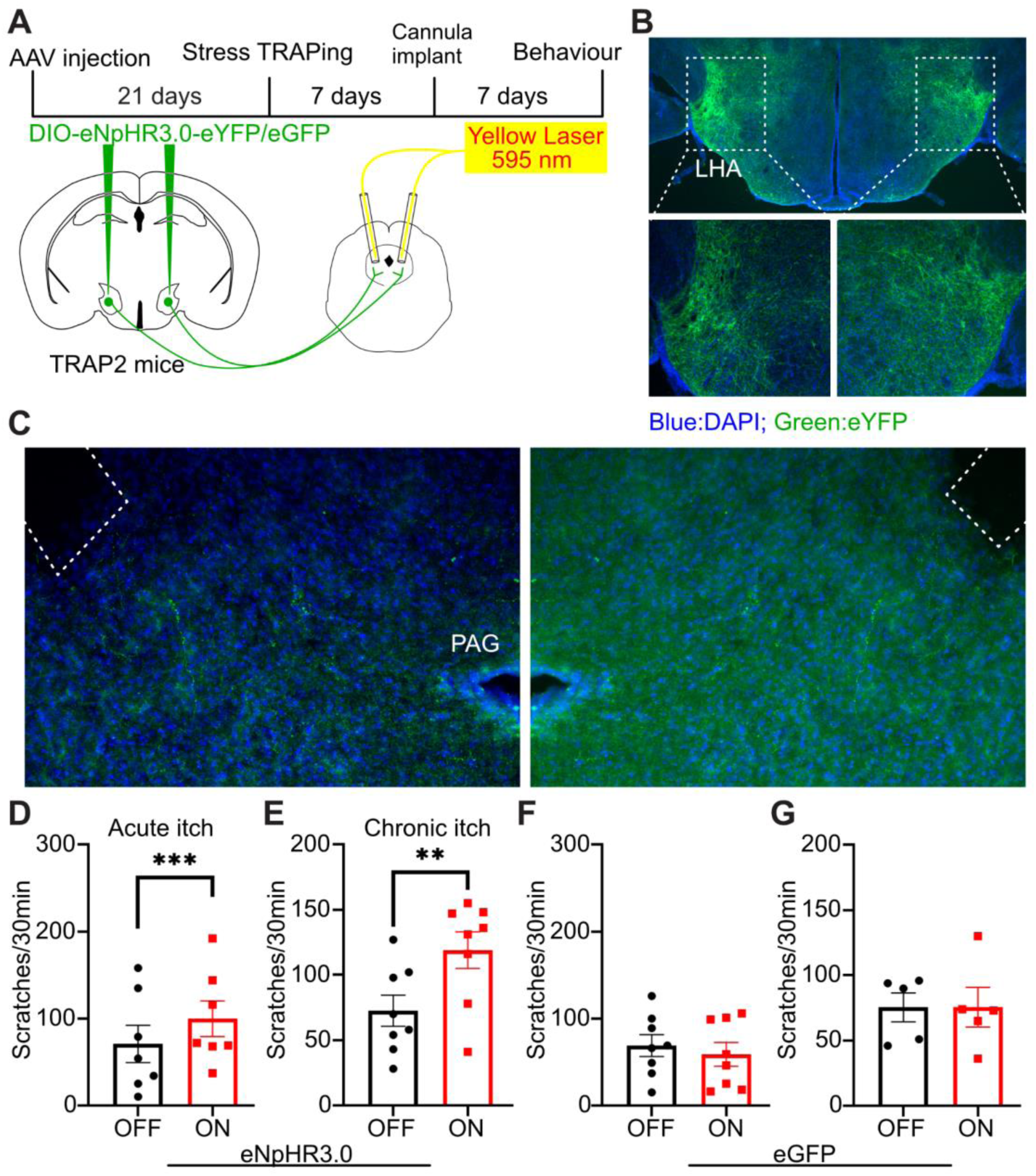
Inhibition of LHA^stress-TRAP^ neurons PAG terminals exacerbate acute and chronic itch. (A) Schematic of the viral injection of eNPHR to inhibit the PAG terminals of the LHA^stress-TRAP^ neurons. (B) Coronal section image of LHA showing the expression of eNPHR in the LHA^stress-TRAP^ neuron, marked by the expression of eGFP (green, blue-DAPI). A higher resolution image of the inset squares is shown at the bottom. (C) Coronal section image of PAG, showing the axon terminals of the LHA^stress-TRAP^ neurons. White dashed lines show fiber tracks. (D) Effect of light ON in PAG (t-test, 29 ± 4.796, *** P = 0.0009, n = 7) on chloroquine-induced scratching compared with light OFF in eNPHR injected mice. (E) Effect of light ON in PAG (t-test, 46.50 ± 10.01, ** P = 0.0024, n = 8) on psoriatic itch induced spontaneous scratching compared with light OFF in eNPHR injected mice. (F) Light ON in PAG had no effect on chloroquine-induced scratching compared with light OFF in eGFP injected mice. (G) Light ON in PAG had no effect on psoriatic itch induced spontaneous scratching compared with light OFF in eGFP injected mice.

## Discussion

The relationship between stress and itch is complex. Here, we shed light on the neural circuitry between the LHA and its downstream neurons in the stress modulation of acute and pathological itch. IEG promoter-mediated gene-trapping strategies allowed us to genetically label restraint stress-sensitive neurons in the LHA. These LHA^stress-TRAP^ neurons were sufficient to suppress acute and chronic itch (Figure 3). Further, these neurons were necessary for acute stress-mediated itch suppression (Figure 3). We mapped the inputs and outputs to find that the LHA^stress-TRAP^ neurons receive somatosensory and affective-motivational inputs from various cortical and subcortical structures and send outputs to them (Figures 4 and 5). Next, the activity of the LHA^stress-TRAP^ neurons corresponded to bouts of spontaneous scratching after mice had developed psoriasis, and can be explained by the altered cellular electrophysiological properties (Figures 6 and 7). We tested through which downstream target LHA^stress-TRAP^ neurons impart their effect on acute and chronic itch. We focused on the brainstem structures PAG, LPBN, and RVM, and found that the LHA-PAG circuitry is sufficient and necessary for the effects of acute RS on itch (Figure 8).

Chronic stress is known to exacerbate both acute and chronic itch, whereas the effects of acute stress on itch remain inconsistent across studies ^55,56^. Importantly, whether neurons in the lateral hypothalamic area (LHA) contribute to these effects has not been addressed. We found that LHA^stress-TRAP^ neurons are mostly glutamatergic (Figure S2). The glutamatergic LHA neurons were activated by stress and shown to be involved in stress modulation of feeding behaviors ^57–59^. Our data shows that chemogenetic activation of LHA^stress-TRAP^ neurons induces anxiety and learned aversion (Figure 2), which is in agreement with previous findings ^60–63^. Acute activation of LHA glutamatergic neurons is aversive in nature, and causes termination of feeding. LHA^stress-TRAP^ neurons mediated anxiety and aversion, likely through their projections to VTA or LHb (Figure 5) ^64,65^. Next, we showed that acute stress or activation of LHA neurons sensitive to acute stress is sufficient for suppressing both the physiological and pathological itch (Figure 3). It remains to be tested if repeated activation of the LHA^stress-TRAP^ neurons is sufficient to cause chronic stress and anxiety, as well as the resultant exacerbation of itch. Interestingly, we find that the LHA^stress-TRAP^ neurons are potentiated by psoriasis (Figure 7).

Inflammatory chronic itch conditions, such as psoriasis, can engage brain areas known to respond to elevated cytokines ^66–69^. In psoriatic mice, where the LHA^stress-TRAP^ neurons start responding to scratching, the information regarding the inflammation leading to psoriasis can likely be transmitted to the LHA through PVN ^70^ (Figure 4), thus increasing the excitability of the LHA^stress-TRAP^ neurons. Alternatively, the sustained activity in the brain nuclei presynaptic to the LHA^stress-TRAP^ neurons that receive somatosensory inputs such as S1, LPBN, and AIC under psoriatic conditions (Figure 4) can potentiate the LHA^stress-TRAP^ neurons. In sum, the LHA^stress-TRAP^ neurons can become sensitive to pruritic stimuli in mice with chronic itch due to increased sustained spontaneous activity in the somatosensory circuits carrying pruritic information or brain circuits engaged by a heightened immune system.

Lateral hypothalamus projects to a wide array of brain regions across the rostro-caudal axis (Figure 5)^12^. Moreover, our data suggest that the stress-sensitive LHA neurons modulate itch via their synaptic connections with the PAG and RVM (Figures 8 and 9). Both of these target nuclei have been shown to modulate itch ^24–28,35,71^. Activation of the inhibitory PAG neural population has been shown to suppress, while inhibition enhances itch ^24,25^. Since LHA^stress-TRAP^ neurons are glutamatergic in nature, it’s likely that these neurons are synapsing on the PAG GABAergic neurons to bidirectionally modulate the physiological and pathological itch. Given the fact that both PAG and RVM terminal activation of the LHA^stress-TRAP^ neurons is able to suppress itch, it will be interesting to test whether the same axon collateral projects to these nuclei to suppress itch or whether two distinct populations of glutamatergic LHA^stress-TRAP^ neurons are involved in itch modulation.

Dysregulation of the HPA axis is a hallmark of psoriasis^72,73^. The affected individuals have difficulties in coping with stress and thus are more prone to anxiety and panic attacks^1^. A subpopulation of neurons in the LHA expressing Orexin has been implicated in regulating corticosterone release through inputs to the PVN^14^. Given the anatomical location of the LHA^stress-TRAP^ neurons and the fact that these neurons are glutamatergic and project to PAG (Figure 5), they likely co-express Orexin^74^. Thus, the LHA^stress-TRAP^ neurons and downstream circuitry can be potentially involved in the stress-coping pathophysiology observed in psoriatic patients^75^.

The caveat of leveraging the Fos-TRAP strategy to study functionally relevant neural populations is that it doesn’t take into account the progress made in dissecting the LHA population according to assigned classes defined by their molecular markers. Glutamatergic Esr1-expressing LHA neurons projecting to the lateral habenula mediate the development of a sex-specific stress state ^76^. While VTA projecting glutamatergic LHA neurons are potentiated by stress, which regulates dopamine release in the prefrontal cortex, thereby promoting stress eating^58^. The midbrain dopaminergic system is known to mediate key aspects of scratch initiation and termination in acute itch^77–80^, thus, LHA-VTA connections can mediate stress-itch interactions. PV+ glutamatergic projection neurons with synaptic connections with PAG in the LHA are nociceptive, and when activated, attenuate acute and persistent pain^81,82^. Orexinergic LHA neurons are involved in the induction and maintenance of negative affective-motivational states such as stress and anxiety^83,84^. Together, molecularly defined excitatory neurons in the LHA with specific axonal targets can explain the effects of LHA^stress-TRAP^ neurons on anxiety levels and itch. In the near future, intersectional genetics ^85^ combined with Fos-TRAP techniques will enable us to combine the molecularly defined neuronal population with the stress-sensitive ones in the LHA and test their roles in stress-modulation of itch and pain. Molecular profiling of the LHA^stress-TRAP^ neurons^86,87^ will shed light on the physiologically relevant neuropeptide or the receptor genes expressed in the cell population of our interest, and may lead to an understanding of the molecular mechanisms underlying stress modulation of pain and itch.

## Abbreviations

4-OHT: 4-hydroxytamoxifen;
AAV: Adeno-associated virus;
ACC: Anterior cingulate cortex;
AcbC: accumbens nucleus cure;
AIC: Anterior insular cortex;
AHC: Anterior Hypothalamic Area central;
AP: anterior-posterior axis;
CeA: Central amygdala nucleus,
BLA: basolateral amygdala;
CM: Central medial thalamus;
CQ: chloroquine;
DCZ: deschloroclozapine;
DREADD: Designer Receptors Exclusively Activated by Designer Drugs;
DTg: Dorsal tegmental nucleus;
eGFP: enhanced Green fluorescence protein;
IL: Infralimbic cortex;
IMQ: Imiquimod;
Irt: Intermediate reticular nucleus;
LAcbSh: Lateral accumbens shell;
LatDN: Lateral dentate nucleus;
LC: locus coeruleus;
LH: Lateral hypothalamus;
LHb: Lateral Hebanula;
LDTg: Laterodorsal tegmental nucleus;
LHA: Lateral Hypothalamic area;
LPBN: lateral parabrachial nucleus;
LPO: Lateral Preoptic Area
LSD: lateral septum nucleus dorsal part;
LSI: lateral septum nucleus intermediate;
LSV: lateral septum nucleus ventral part;
MPA: Medial preoptic area;
mPBN: medial parabrachial nucleus;
mRt: mesencephalic reticular formation;
MO: Medial orbital cortex;
MS: Medial Septum;
PAG: Periaqueductal gray;
PLH: peduncular part lateral hypothalamus;
PO: Preoptic area;
PrL: Prelimbic cortex;
PV: paraventricular thalamus;
PVN: Paraventricular hypothalamus nucleus;
Re: thalamic reuniens nucleus;
RSGc: Retrosplenial granular cortex;
RVM: rostral ventromedial medulla;
S1: Primary somatosensory cortex;
SC: Superior Colliculus;
SNc: Substantia Nigra pars compacta;
*Slc17a6*: Solute carrier family 17 member 6 (Vglut2- Vesicular glutamate transporter);
*Slc32a1*: Solute carrier family 32 member 1(GABA vesicular transporter)
SoI: solitary nucleus;
STLP: Bed nucleus of stria terminalis, lateral division, posterior;
STLV: Bed nucleus of stria terminalis, lateral division, ventral;
TRAP2: Targeted Recombination in Active Populations;
VDB: nucleus of the vertical limb of the diagonal band;
VTA: Ventral Tegmental area.

---

**Figure S1:**
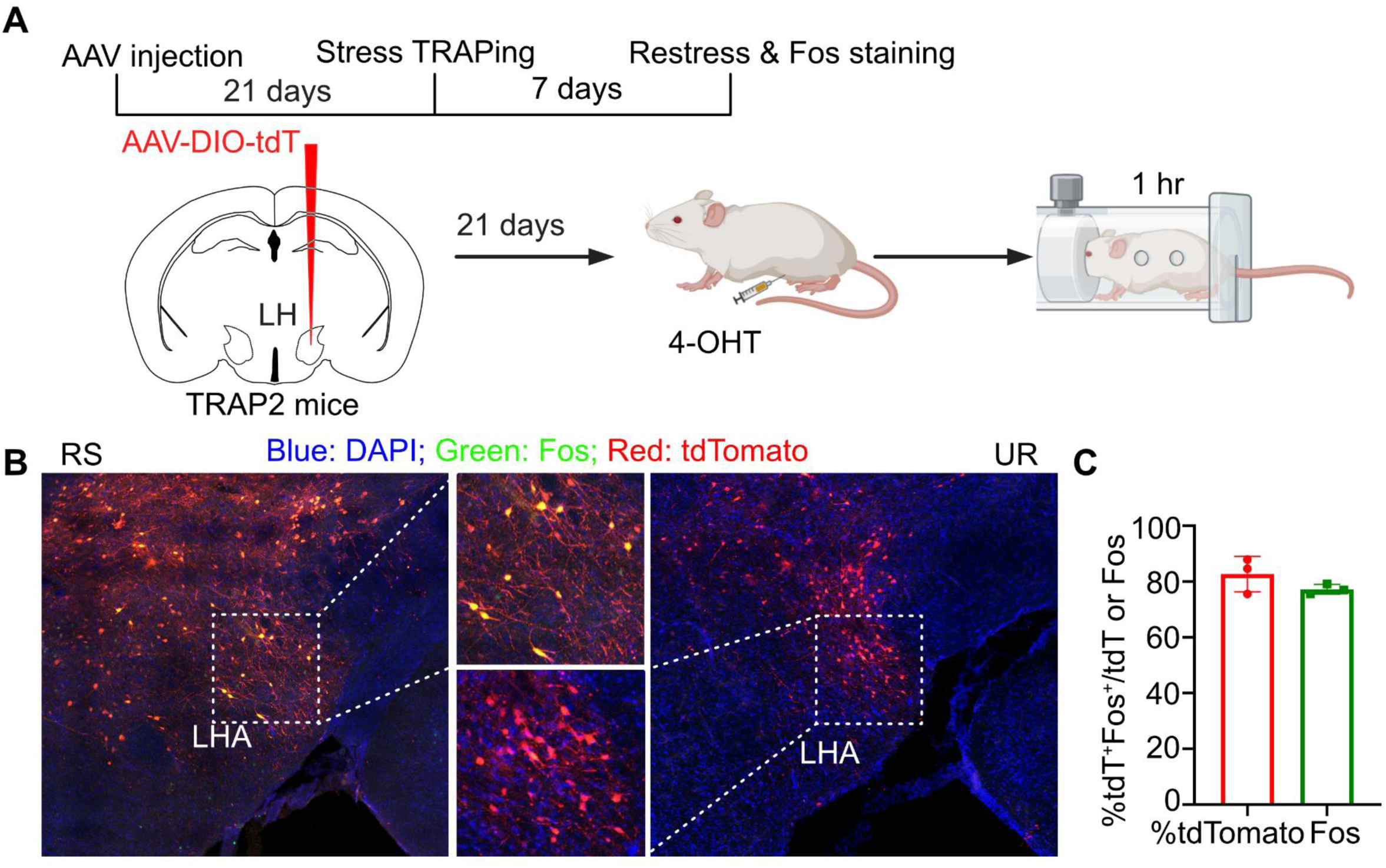
TRAPing of LHA stress-sensitive neurons. (A) Schematic of the experiment timeline. AAV encoding Cre-dependent tdTomato was injected into the LHA of the TRAP2 mice. (B) Coronal section confocal image of the LHA^stress-TRAP^ neurons shows the overlap between tdTomato (red) positive cells and Fos (green) positive cells. The middle image shows the zoomed-in image from the marked squares. Yellow cells are both positive. (C) Quantification of B. The red bar indicates the percentage of tdTomato-positive cells also expressing Fos (mean ± SD: 77.29 ± 1.78), and the green bar represents the percentage of Fos-positive cells also expressing tdTomato (mean ± SD: 82.75 ± 6.37) from n = 3 mice, six sections per mouse.

**Figure S2:**
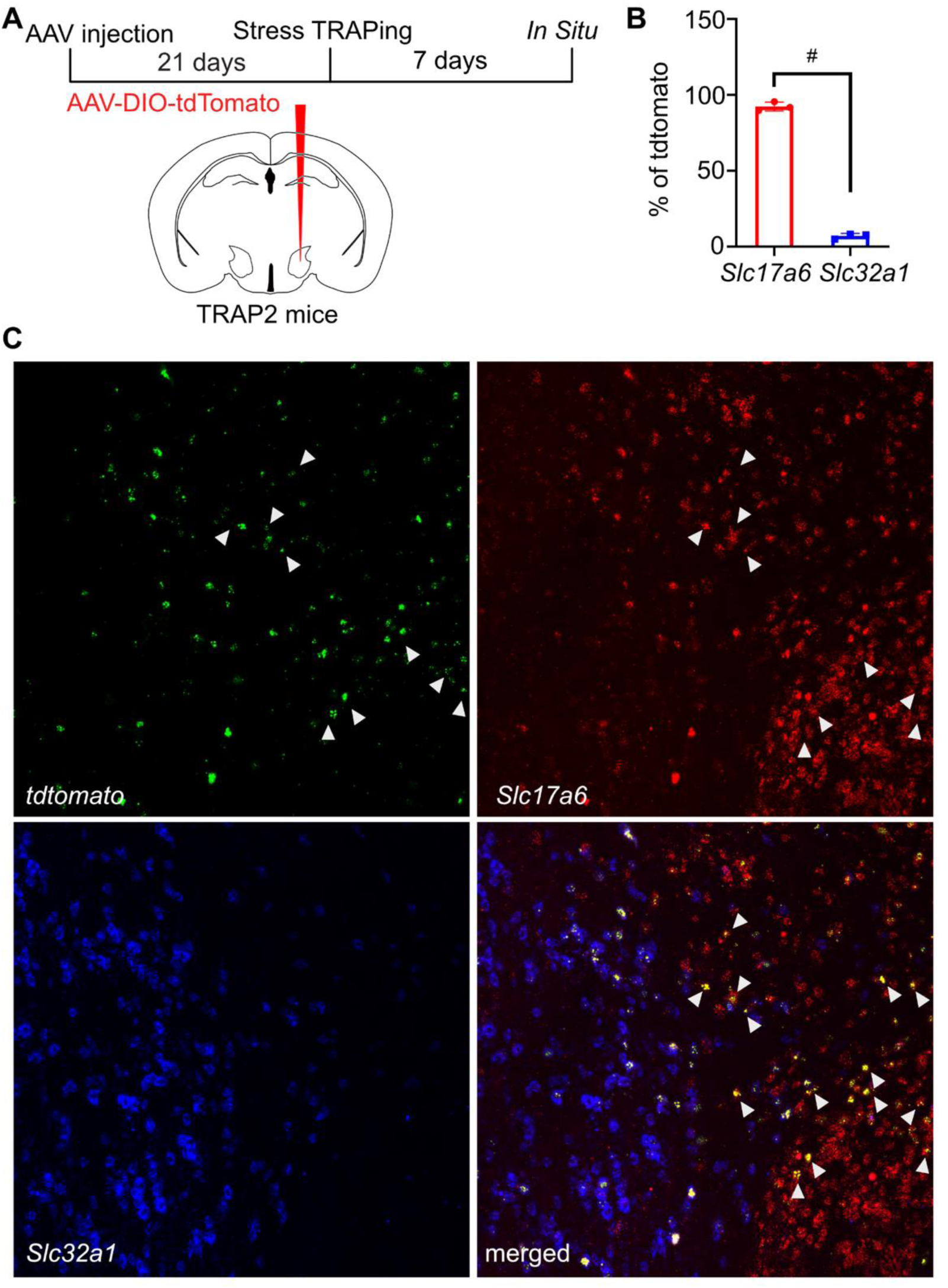
LHA^stress-TRAP^ neurons are mostly glutamatergic neurons. (A) Schematic of the experiment timeline. AAV encoding Cre-dependent tdTomato was injected into the LHA of the TRAP2 mice to mark the LHA^stress-TRAP^ neurons with tdTomato expression. (B) Quantification of the overlap of tdTomato with *Slc17a6* and *Slc32a1*. % of tdTomato overlapping with Slc17a6 is 92.38 ± 0.79 and the % of tdTomato overlapping with *Slc32a1* is 6.9 ± 0.47. Data is represented as mean ± SEM, n=3. Means difference in overlap is, unpaired t-test, 85.49 ± 1.96, # P <0.0001, n = 3. (C) Multiplexed *in situ* hybridization for *tdTomato* (green), *Slc17a6* (red), *Slc32a1* (blue). Bottom right shows the merged image. The white triangles show the overlap (yellow) between the *tdTomato* and the *Slc17a6*.

**Figure S3:**
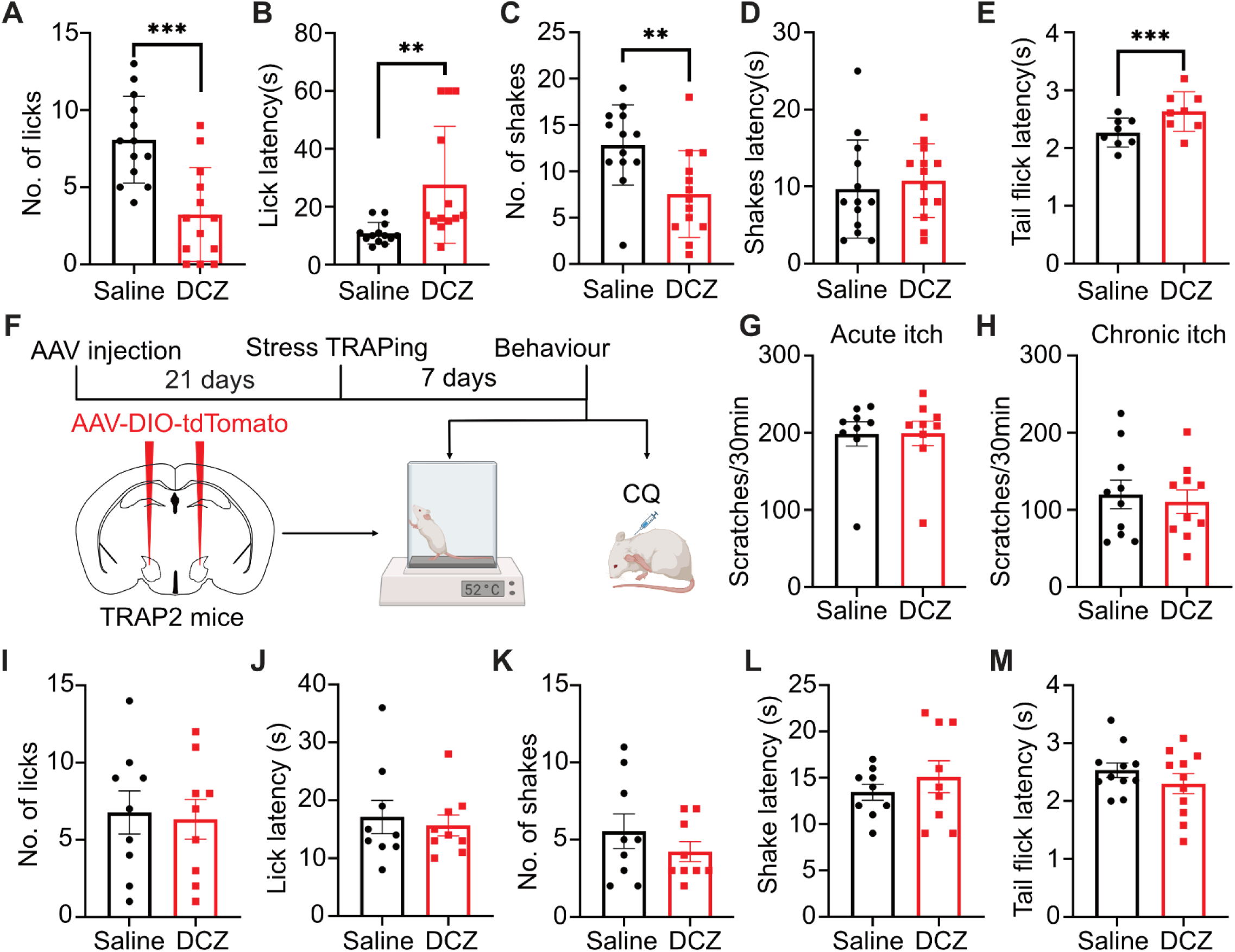
LHA^stress-TRAP^ neuron activation suppresses thermal pain. (A) Effect of i.p. DCZ (t-test, 4.846 ± 0.966, ***P = 0.0003, n = 13) administration on the number of licks compared with i.p. saline on a hotplate set at 52°C. (B) Effect of i.p. DCZ (t-test, 16.77 ± 5.372, **P = 0.0088, n = 13) administration on lick latency compared with i.p. saline on a hotplate set at 52°C. (C) Effect of i.p. DCZ (t-test, 5.308 ± 1.5, **P = 0.0041, n = 13) administration on the number of shakes compared with i.p. saline on a hotplate set at 52°C. (D) Effect of i.p. DCZ (t-test, 1.077 ± 1.998, ns P = 0.599, n = 13) administration on shake latency compared with i.p. saline on a hotplate set at 52°C. (E) Effect of i.p. DCZ (t-test, 0.3638 ± 0.0613, ***P = 0.0006, n = 8) administration on tail flick latency compared with i.p. saline. (F) Schematic of the experiment. AAV encoding Cre-dependent tdTomato was bilaterally injected into the LHA of the TRAP2 mice. (G) DCZ administration had no effect on chloroquine induced scratching compared with i.p. Saline (controls) in tdTomato injected mice. (H) DCZ administration had no effect on psoriatic itch-induced spontaneous scratching in tdTomato injected mice. (I) DCZ administration had no effect on the number of licks on a hotplate set at 52°C in tdTomato injected mice. (J) DCZ administration had no effect on lick latency on a hotplate set at 52°C in tdTomato injected mice. (K) DCZ administration had no effect on the number of shakes on a hotplate set at 52°C in tdTomato injected mice. (L) DCZ administration had no effect on shake latency on a hotplate set at 52°C in tdTomato injected mice. (M) DCZ administration had no effect on tail flick latency on a hotplate set at 52°C in tdTomato injected mice.

**Figure S4:**
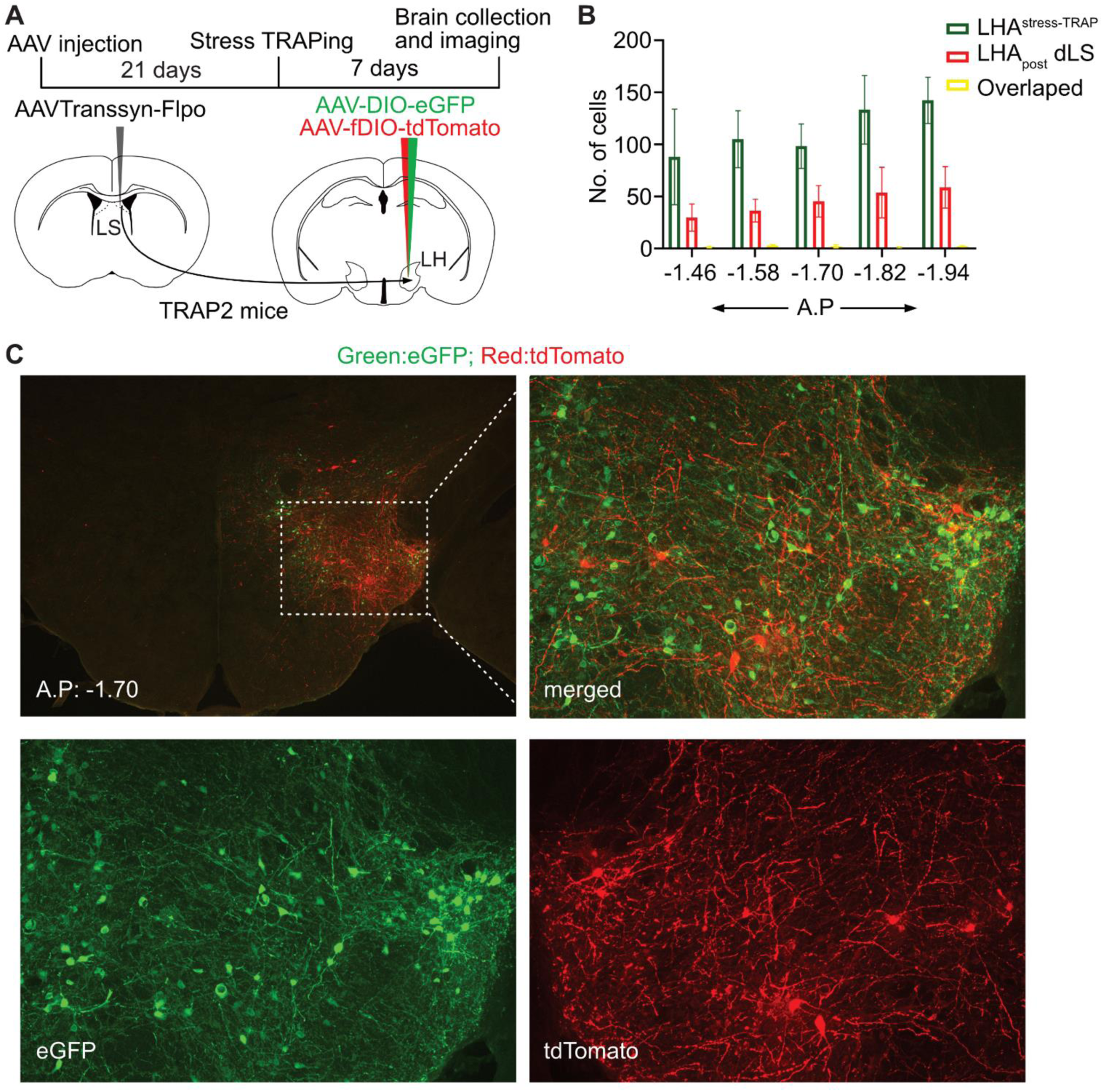
LHA^stress-TRAP^ neurons are distinct from the LHA_post-LS_ neurons. (A) Schematics of the experiment. Transsynaptic Flpo encoding AAV was injected into the LS of the TRAP2 mice. Two AAVs encoding Cre-dependent GFP and Flpo dependent tdTomato were mixed in 1:1 ratio and injected into the LHA of the same mice. (B) Quantification of eGFP expressing (LHA^stress-TRAP^) neurons and tdTomato expressing (LHA_post-LS_) neurons across anterior-posterior axis. (C) Coronal section of LHA showing the expression of eGFP and tdTomato. Higher resolution image of the inset square is shown on the right.

**Figure S5:**
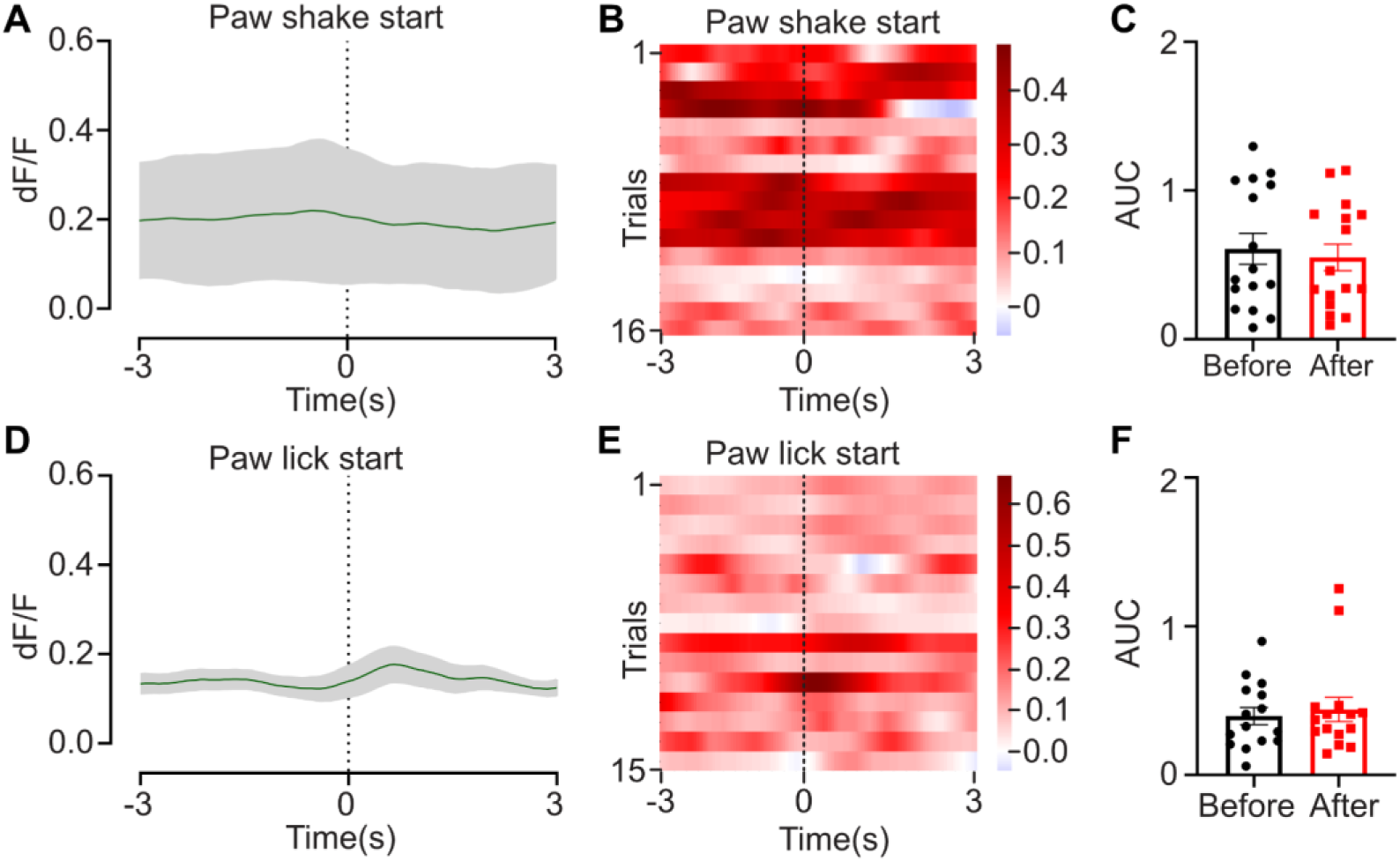
LHA^stress-TRAP^ neurons does not respond to nocifensive behaviours on the hotplate test. (A) The average fluorescent signal at the start of hind paw shake when mice were challenged on the hotplate test (B) Heatmap showing the response during the hind paw shake. (C) Area under the curve (AUC) 3s before and after the start of paw shake. (D) The average fluorescent signal at the start of hind paw lick when mice were challenged on the hotplate test (E) Heatmap showing the response during the hind paw lick. (F) Area under the curve (AUC) 3s before and after the start of paw lick.

**Figure S6:**
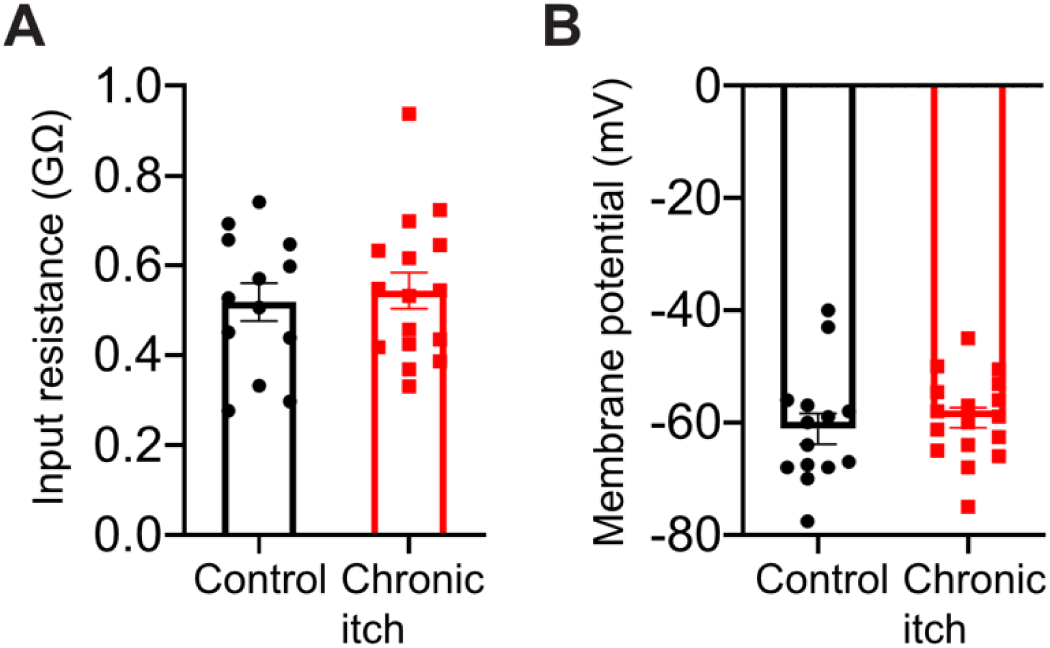
Activation of RVM terminals of the LHA^stress-TRAP^ neuron suppresses thermal. (A) Schematic of the experiment timeline. AAV encoding Cre-dependent hM3Dq was injected bilaterally into the LHA of the TRAP2 mice. One week after the stress-TRAPing, cannula was implanted at the RVM. (B) Schematic of the DCZ/saline infusion at the RVM. (C) Effect of DCZ infusion at the RVM (t-test, 2.286 ± 0.8371, *P = 0.0342, n = 7) on the number of licks on a hotplate set at 52°C. (D) Effect of DCZ infusion at the RVM (t-test, 14.43 ± 4.303, *P = 0.0154, n = 7) on lick latency compared to the control condition on a hotplate set at 52°C. (E) Effect of DCZ infusion at the RVM (t-test, 7 ± 1.38, **P = 0.0023, n = 7) on the number of shakes compared with saline infusion on a hotplate set at 52°C. (F) Effect of DCZ infusion at the RVM (t-test, 18 ± 3.032, **P = 0.001, n = 7) on shake latency on a hotplate set at 52°C. (G) Effect of DCZ infusion at the RVM (t-test, 0.6129 ± 0.20, *P = 0.023, n = 7) on tail flick latency compared with saline infusion. (H) DCZ infusion at RVM had no effect on the number of licks compared with saline infusion on a hotplate set at 52°C in the tdTomato injected mice. (I) DCZ infusion at RVM had no effect on lick latency compared with saline infusion on a hotplate set at 52°C in the tdTomato injected mice. (J) DCZ infusion at RVM had no effect on the number of shakes on a hotplate set at 52°C in the tdTomato injected mice. (K) DCZ infusion at RVM had no effect on shake latency on a hotplate set at 52°C in the tdTomato injected mice. (L) DCZ infusion at RVM had no effect on tail flick latency on a hotplate set at 52°C in the tdTomato injected mice.

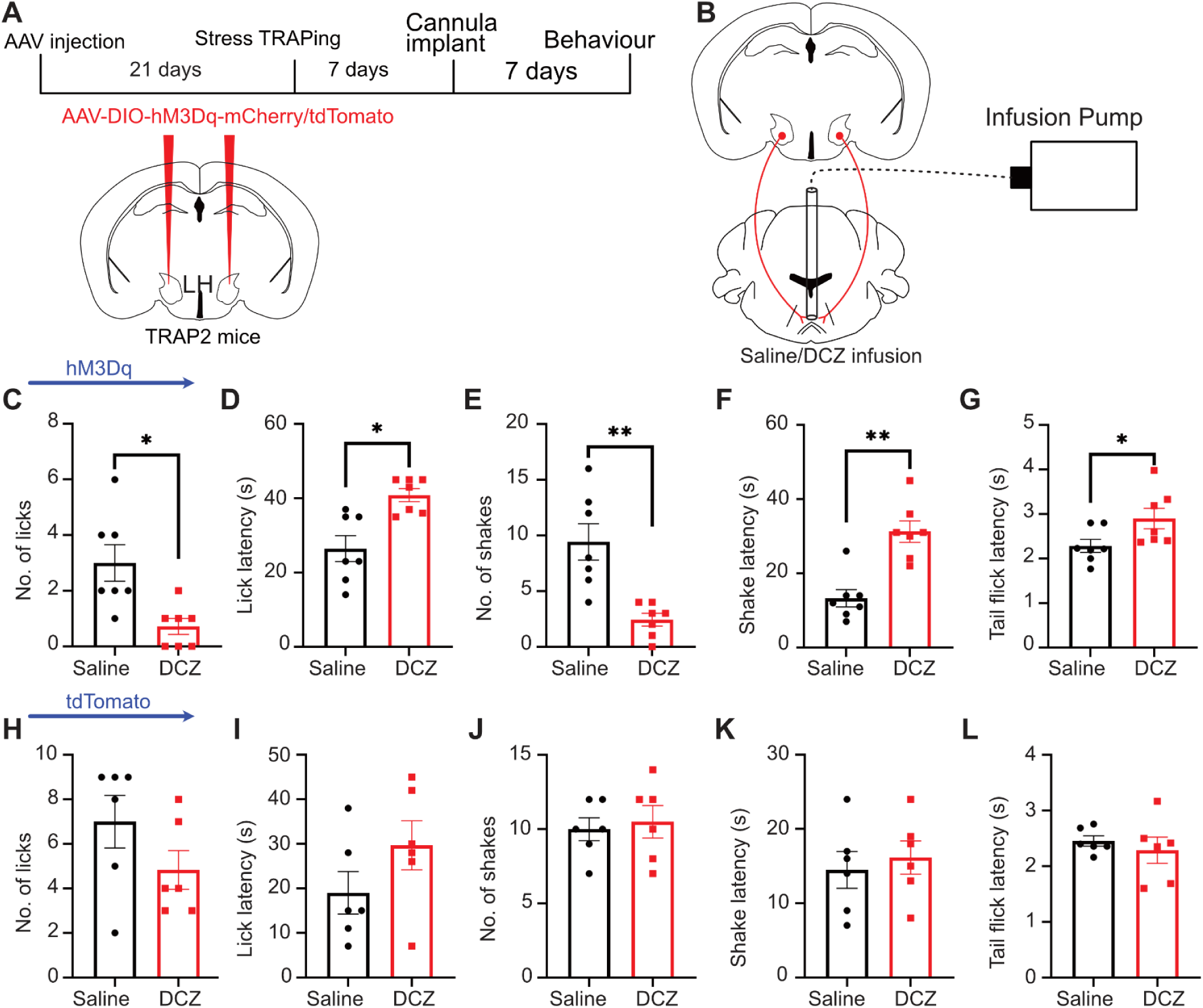

## Authorship contributions

J.N.P., A.H., M.P. performed experiments, J.N.P., A.H., M.P. analyzed data, J.N.P., A.H., G.S., and A.B. conceptualized and designed the study, and wrote the manuscript, G.S. and A.B. obtained funding.

